# Pressorum Sensing: Growth-induced Compression Activates cAMP Signaling in *Pseudomonas aeruginosa*

**DOI:** 10.1101/2024.07.08.602437

**Authors:** Lei Ni, Yajia Huang, Yaoxin Huang, Yue Yu, Jiarui Xiong, Hui Wen, Wenwen Xiao, Haiyi Liang, Fan Jin

## Abstract

Bacteria employ various strategies to coordinate population-level behaviors, with quorum sensing being a well-established mechanism. Here, we report a novel population-level regulatory mechanism in *Pseudomonas aeruginosa*, which we term ‘pressorum sensing’. This mechanism allows bacteria to modulate their collective behavior in response to growth-induced mechanical compression in confined spaces. Using a highly sensitive cAMP biosensor in combination with microfluidics, we demonstrate that when compressive forces reach approximately 30 nN, *P. aeruginosa* cells rapidly increases intracellular cAMP levels via the Pil-Chp chemosensory system. This response leads to up-regulation of the Type III Secretion System, a key virulence factor. Unlike quorum sensing, which relies on diffusible chemical signals, pressorum sensing utilizes mechanical cues to gauge population density and spatial confinement. In bacterial colonies, this mechanism generates striking spatial patterns of cAMP signaling, including traveling rings that coincide with step-like structures in colony morphology. Our findings reveal a previously unknown link between mechanical compression and bacterial virulence, providing new insights into how *P. aeruginosa* coordinates population-level responses in confined environments. This work also expands our knowledge of mechanogenetics and opens up new possibilities in synthetic biology and bioengineering applications.

## Introduction

Microorganisms have evolved sophisticated mechanisms to sense and respond to various environmental cues, enabling them to adapt to different stages of life cycle and optimize their survival strategies. Among these, quorum sensing is a well-studied phenomenon that allows bacteria to monitor their population density through the secretion and detection of chemical signaling molecules called autoinducers ^1,2^. This mechanism enables bacteria to coordinate their gene expression at the population level, leading to significant biological consequences, such as the regulation of virulence factors and the formation of biofilms.

The triggering of quorum sensing requires a relatively high local cell density, but it does not necessitate the bacteria to enter an extremely crowded state of mutual contact, as the process can be initiated in a liquid medium. However, bacteria in many natural habitats, such as biofilms, tissue crevices, and the confinement inside phagosomes, often grow in densely packed environments where they experience mechanical compression ^3–7^. The impact of physical confinement on bacterial behavior and its implications for population-level regulation remain largely unexplored. Serendipitously, while investigating changes in the second messenger cAMP in *Pseudomonas aeruginosa*, we observed an intriguing phenomenon: when bacteria grow in confined spaces, transitioning from sparse to densely packed states, their intracellular cAMP levels rise significantly. We later demonstrated that densely packed growth can trigger the expression of cAMP downstream genes related to the type III secretion system.

In *P. aeruginosa*, cAMP is a crucial second messenger that regulates the expression of numerous virulence factors, including type II and III secretion systems and type IV pili ^8–10^. Previous studies have demonstrated that surface contact can trigger cAMP signaling through the extension and retraction of type IV pili ^11,12^. Our findings build upon this knowledge, revealing that bacteria can further augment their cAMP signaling responses when exposed to an additional mechanical cue-compression in densely packed growth conditions - following initial surface colonization. Subsequent experiments confirmed that this response is independent of extracellular type IV pili and instead depends on the extent of mechanical compression generated by their own growth, a mechanism that bears resemblance to how they detect their population size through chemical signals in quorum sensing systems ^13,14^. Both of the two sensory mechanisms allow bacteria to modulate population-level phenotypic changes in response to increasing population size. Drawing inspiration from the concept of quorum sensing, we propose the term ‘pressorum sensing’ to describe this novel mechanism.

The identification of pressorum sensing expands our understanding of how bacteria adapt to and thrive in densely packed environments, and we showed that virulence gene expression is also subject to population size control. This finding complements the well-established concept of quorum sensing and provides new insights into bacterial communication and adaptation strategies in crowded habitats.

## Results

### Growth-induced compression in confined space triggers a Pil-Chp-mediated cAMP response in *P. aeruginosa*

To investigate the rapid temporal dynamics of cAMP fluctuations within *P. aeruginosa* cells, we employed Gflamp1, a highly sensitive and specific monomeric circularly permuted fluorescent protein (cpFP) sensor ^15^. With a dissociation constant of 2 µM, Gflamp1 aligns with the physiological range of cAMP concentration in *P. aeruginosa*. We fused Gflamp1 to CyOFP1, a fluorescent protein with constant fluorescence independent of cAMP concentration (Fig. 1A) ^16^, enabling normalization of the Gflamp1 signal for variations in cell density and colony thickness. Gflamp1 and CyOFP1 share an excitation wavelength of 488 nm, allowing simultaneous excitation by a single laser beam, while their distinct emission spectra facilitate separation and quantification of their respective fluorescence signals. The Gflamp1/CyOFP1 ratio serves as a reliable readout for cAMP concentration, independent of excitation laser intensity and corrected for variations in cell density and colony thickness. This ratiometric approach, combined with dual-view imaging, enables accurate, high spatiotemporal resolution measurements of cAMP levels in living cells, even in densely packed environments such as biofilms.

**Figure 1:**
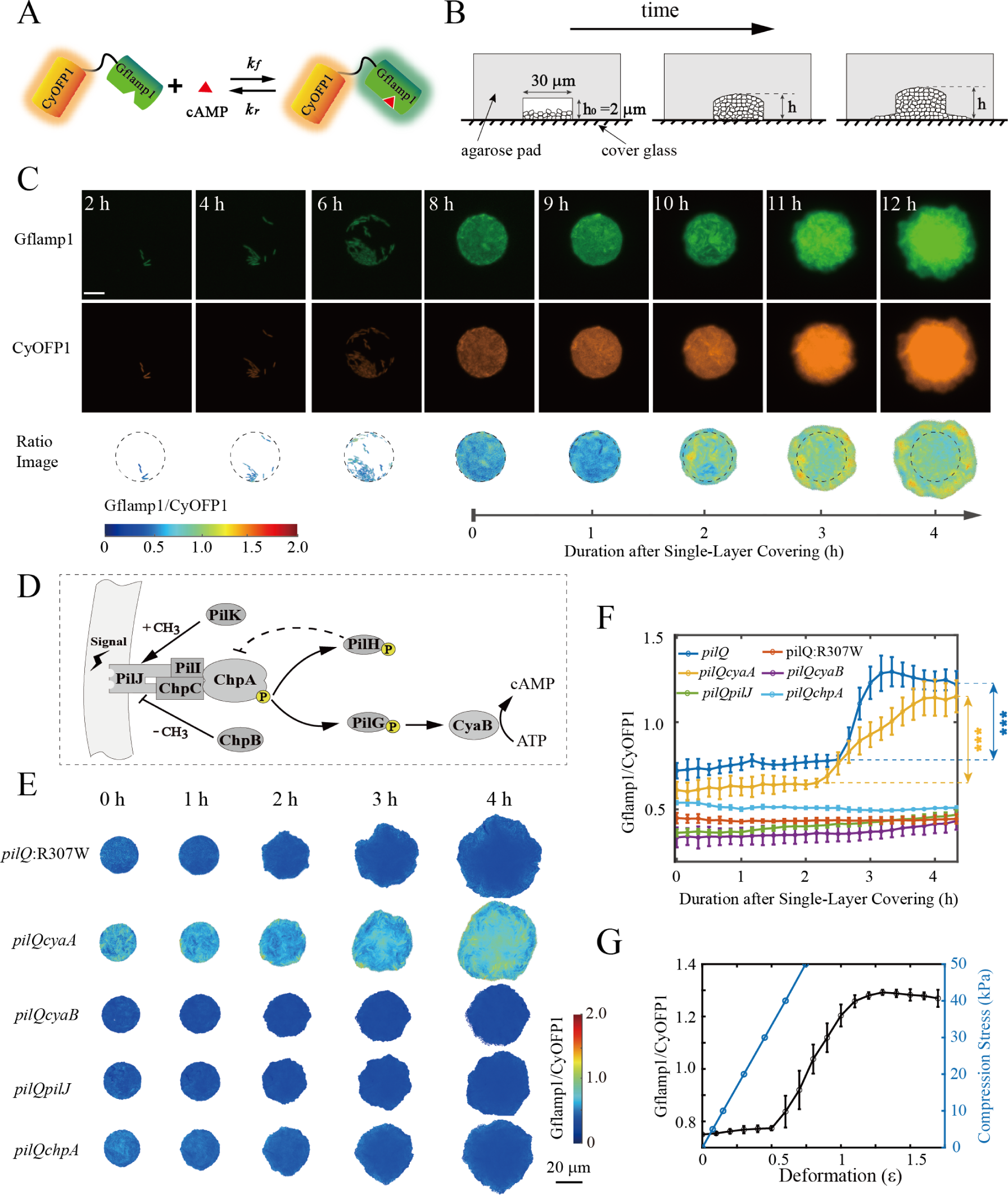
Confined growth-induced compression triggers a Pil-Chp-mediated cAMP response in *P. aeruginosa*. (A) Schematic of the Gflamp1-CyOFP1 cAMP biosensor. (B) Time-lapse images of *P. aeruginosa* growth in agarose microcavities (30 µm diameter,2 µm depth). Scale bar: 10 µm. (C) Gflamp1 and CyOFP1 fluorescence, ratio images, and quantification of the Gflamp1/CyOFP1 ratio during single-layer to multi-layer transition, the dashed lines indicate the boundaries of the microcavity. (D) Schematic of the Pil-Chp chemosensory system regulating CyaB activity and cAMP synthesis. (E) Time-resolved fluorescence ratio images of *pilQ* mutant strain and additional gene mutant strains derived from the *pilQ* mutant background. Scale bar: 20 µm. (F) Quantification of the Gflamp1/CyOFP1 ratio in *P. aeruginosa* mutant strains. Statistical significance determined by t-test, ∗∗∗, *p <* 0.001. (G) Relationship between gel deformation *ε*, compressive pressure, and the Gflamp1/CyOFP1 ratio.

Building upon our initial observations(see Supplementary Information), we hypothesized that the mechanical compression experienced by *P. aeruginosa* cells during the self-driven jamming state could trigger an elevation in their intracellular cAMP levels. This hypothesis is supported by the synchronized increase in the Gflamp1/CyOFP1 ratio observed as the cells transitioned from a fluidic-like motility to a densely packed, jammed state (Fig. S1A, S1B, movie S1: wild type (1.5%)). Furthermore, the higher cAMP levels detected at the expanding periphery of the twitching colony (Fig.S1A) suggest that cells experiencing greater mechanical compression, possibly due to the resistance from the agarose matrix, exhibit elevated cAMP signaling. To fully understand the mechanisms underlying the increased intracellular cAMP levels following bacterial jamming, several additional factors must be considered, including the potential accumulation of signaling molecules due to high cell density, modulation of type IV pili dynamics during the motility state transition, and the impact of nutrient deprivation on cAMP signaling.

To investigate whether the accumulation of species-specific signaling molecules or the dynamics of type IV pili-mediated surface motility contribute to the synchronized elevation of cAMP levels, we co-cultured *P. aeruginosa* with *Escherichia coli* at a 1:10 ratio beneath agarose pads. Again, we observed an increase in the Gflamp1/CyOFP1 signal within *P. aeruginosa* cells after cells became densely packed (Fig. S2, Movie S2), ruling out the potential induction of cAMP increase by accumulated *P. aeruginosa*-specific secreted factors. To further examine the role of type IV pili in the cAMP response, we co-cultured *E. coli* cells with *pilQ* mutant cells of *P. aeruginosa*, which lacks functional surface pili (Fig. S3) and twitching motility ^17^. Remarkably, we still detected a similar increase in the Gflamp1/CyOFP1 signal in the *pilQ* mutant under densely-packed conditions (Fig. S2, Movie S2), indicating that surface pili dynamics are not a determining factor for the intracellular cAMP response. Twitching motility mediates collective migration of cells beneath the agarose pad ^18,19^, leading to constantly changing compaction states within the population, and pilus retraction could potentially modulate cAMP signaling ^11,12^. Therefore, we chose the non-piliated *pilQ* mutant for subsequent experiments to rule out the influence of twitching motility and surface type IV pili appendages on the synchronized elevation of cAMP levels, as it serves as a better genetic background compared to the *pilA* mutant, which can still influence intracellular signaling through the PilSR and PilJ-Chp system ^11,20^.

To better control for nutrient or oxygen depletion and establish a more robust experimental system, we modified the agarose pad assay by creating cylindrical microcavities on the gel surface with diameters of 30 µm and 60 µm and a depth of 2 µm (Fig. 1B). These microcavities, covered with a coverslip, formed enclosed chambers wherein growing bacteria could enter a densely packed state earlier upon pushing against the chamber wall (Fig. 1B), while the shallow depth ensured sufficient nutrient availability during the crowding process. Cells initially grew as a monolayer with constant CyOFP1 fluorescence. After forming a complete monolayer, the CyOFP1 signal increased, indicating vertical multilayer formation, but the Gflamp1/CyOFP1 ratio remained unchanged (Fig. S4). Approximately two hours later, when the cells had filled the microcavities, the excessive growth pressure caused delamination between the agarose pad and the coverslip at the edge of cavities, allowing bacteria to leak out and grow outwards (Fig. 1C). It was at this time that the Gflamp1/CyOFP1 ratio significantly increased from 0.7 to 1.3 (*p <* 0.001) and reached a plateau (Fig. 1F, Fig. S4 and Fig. S5), a phenomenon observed in both cavity sizes. This observation strongly suggests that the signal elevation is triggered by the compression forces. Replacing Gflamp1 with the cAMP-binding deficient mutant Gflamp1:R307W ^21^ abolished the fluorescence ratio change (Fig. 1E, 1F), confirming the specificity of the cAMP response. These results, combined with our previous findings, led us to conclude that densely-packed crowded growth triggers an intracellular cAMP response mechanism in *P. aeruginosa*, with mechanical compression potentially being the upstream signal. The use of microcavities with defined geometries provides a well-controlled experimental system to study this phenomenon while minimizing confounding factors.

Next, we investigated the intracellular signaling pathways underlying the observed cAMP response. *P. aeruginosa* has two adenylate cyclases, CyaA and CyaB ^9^, responsible for cAMP synthesis, and a single phosphodiesterase CpdA that catalyzes cAMP degradation ^22^. Through gene knockout experiments, we determined that the heightened cAMP response during densely packed growth was abolished solely in the *cyaB* mutant strain (Fig. 1E, 1F). CyaB activity is regulated by the Pil-Chp chemosensory system ^23^, which consists of the transmembrane chemoreceptor PilJ, the histidine kinase ChpA, and the response regulators PilG and PilH (Fig. 1D). The cAMP response was abrogated in the *pilJ* and *chpA* mutant strains(Fig. 1E, 1F), indicating that the Pil-Chp system mediates the cAMP response during densely packed growth. To estimate the compressive forces acting on bacteria when the Gflamp1/CyOFP1 signal responds, we established a simplified model based on cell deformation (see Supplementary Information). Using CyOFP1 intensity to estimate the number of cell layers within the chamber, we calculated the apparent strain of bacterial cells (*ε* = (*h − h*_0_)/*h*_0_ = *ε_e_* + *ε_g_*) and generated a relationship curve between *ε* and Gflamp1/CyOFP1. The transition point for cAMP response occurred when *ε* was about 0.5, corresponding to a compressive pressure of ∼30 kPa (Fig. 1G, see Supplementary Information), translating to a compressive force of ∼30 nN acting on individual cells.These findings demonstrate that the Pil-Chp chemosensory system, through the regulation of CyaB activity, mediates the cAMP response triggered by mechanical compression during densely packed bacterial growth.

### Activation of Type III Secretion System Gene Expression During Densely-packed Bacterial Growth

One question is whether the intracellular cAMP response caused by growth-induced compression indicates a change in bacterial phenotype. In *P. aeruginosa*, cAMP forms a complex with the transcription factor Vfr to regulate the expression of multiple genes ^24^, among which the Type III secretion system (T3SS) genes have garnered significant interest. The cAMP-Vfr complex directly regulate the transcription of *exsA* ^10^, which encodes the master regulator ExsA for T3SS gene expression ^25^ (Fig. 2A). T3SS is a critical tool for *P. aeruginosa* to infect host cells, and its activation is a characteristic bacterial phenotype during acute infection ^26^. To examine the expression of T3SS genes during densely-packed bacterial growth, we constructed transcriptional reporters with *sfGFP* and *cyOFP1* genes expressed downstream of a seleted T3SS promoter and a constitutive promoter, respectively. CyOFP1 fluorescence was used to normalize the variation of plasmid copy number and cell density, promoter activities were represented by SfGFP/CyOFP1.

**Figure 2:**
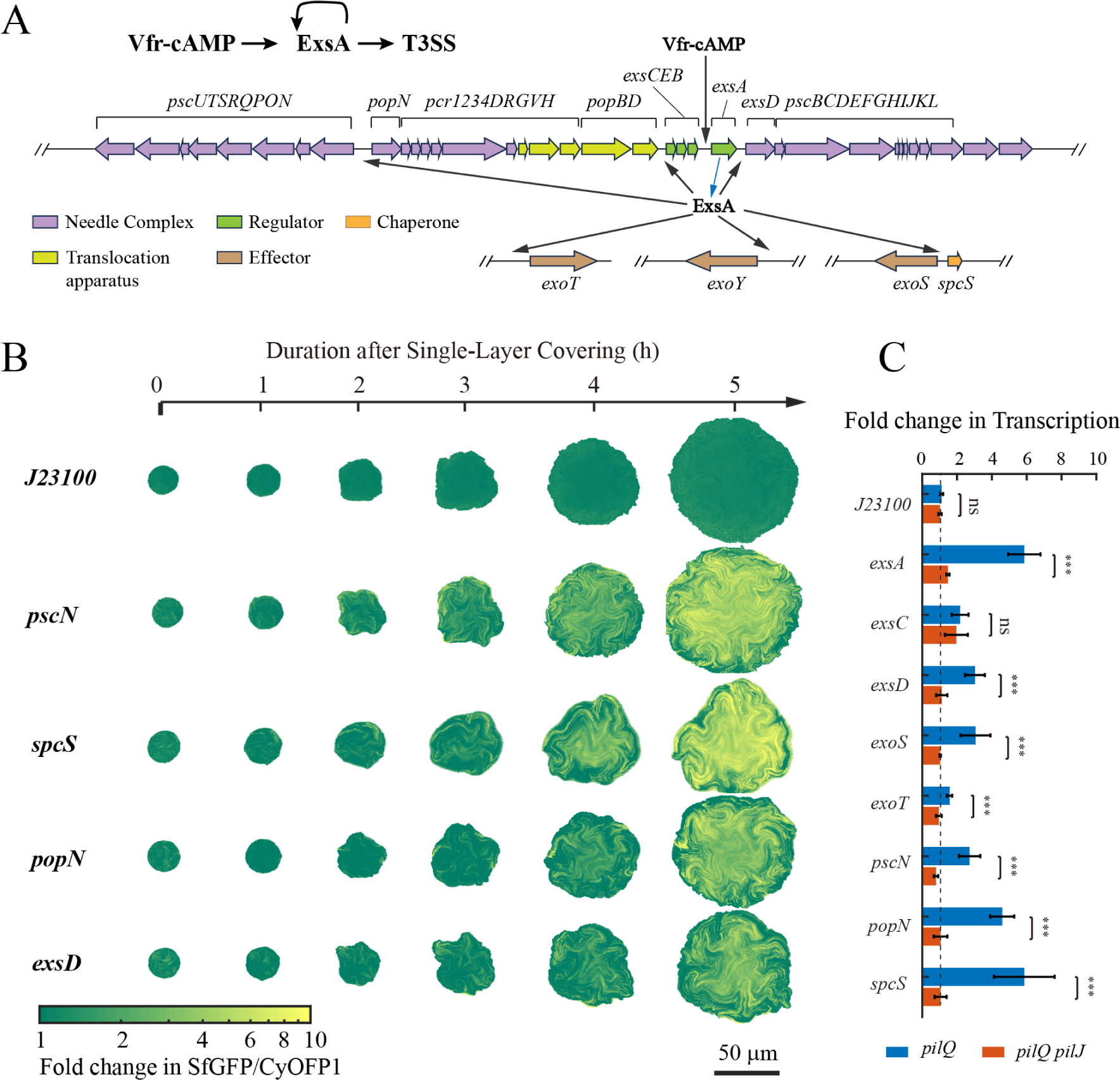
Confined growth induces the expression of T3SS promoters in *P. aeruginosa*. (A), Illustration depicting cAMP-Vfr complex regulation of type III secretion system genes via ExsA. (B) Time-series images of T3SS transcriptional reporters (expressed in *pilQ* mutant strain) growing in agarose microcavities (30 µm diameter, 2 µm depth). Colormap indicates promoter expression levels normalized to the SfGFP/CyOFP1 ratio at the point when chambers were just single-layer covered. Scale bar: 50 µm. (C) Fold change in T3SS promoter expression, comparing levels at 5 generations post-monolayer formation to those at initial monolayer establishment, in *pilQ* and *pilQpilJ* mutant strains. Statistical significance determined by t-test, *^∗∗∗^*, *p <* 0.001, ns, non significant.

Promoter activities, normalized by SfGFP/CyOFP1 signals at the onset of single-layer coverage, were visualized using colormaps (Fig. 2B). The majority of T3SS promoters exhibited increased activity 3-4 hours post single-layer formation (Fig. 2B, Movie S4), corresponding to 1-2 bacterial growth cycles after the cAMP signaling response. Five hours after single-layer coverage, a 3-6 fold increase in average SfGFP/CyOFP1 was observed, while the constitutive promoter *J23100* remained unaltered(Fig. 2C). Repeating the measurements in the *pilJ* mutant strain revealed no change in T3SS promoter activities (Fig. 2C). These findings collectively indicate a global upregulation of T3SS gene expression during dense bacterial growth, subsequent to the mechanically-induced cAMP signaling.

### Introduction of ‘Pressorum Sensing’

Given the observed phenomenon of microorganisms exhibiting elevated intracellular cAMP concentration and virulence phenotype switching in response to mechanical compression, we introduce the term ‘pressorum sensing’ to describe this behavior. ‘Pressorum sensing’ is an analogicaconcept of ‘quorum sensing’, reflecting a group of bacterial cells change their physiological state when the local mechanical pressure originated from intercellular compression exceeds a certain threshold. The ‘pressorum sensing’ system utilizes mechanical force as the transduced signal, not the chemical signals employed in ‘quorum sensing’. However, both systems share a common feature to enable the detection of population size. Quorum sensing measures population size through the equilibrium between self-secretion and diffusion of autoinducers, reflecting the chemical dimension of population growth. In contrast, pressorum sensing measures population size through the balance between the growth-dependent expansion pressure and release of it by environmental physical boundaries, reflecting the physical dimension of population growth. The term ‘pressorum’ is derived from ‘com**press**ion’ and ‘qu**orum**’. This newly coined term captures the unique ability of microorganisms to sense and respond to mechanical compression during crowded growth in confined habitats.

### Direct Mechanical Compression at Single-cell Level Activates cAMP Signaling

In quorum sensing, the expression of downstream genes can be activated in a single bacterium if the autoinducer concentration is sufficient^27^. We wonder whether pressorum sensing shares similar characteristics that the cAMP signaling response could be triggered by applying mechanical pressure to individual bacteria, independent of cell-cell contact. To directly apply mechanical pressure to single bacteria, we designed a bilayer microfluidic device (Fig. 3A shows a schematic diagram; see Fig. S6 for detailed device configuration). The device consists of two chambers separated by a 50-µm-thick PDMS membrane with cylindrical protrusions on its underside. The membrane and the upper PDMS chip form an closed chamber, which is filled with PBS buffer and connected to an air pressure pump. The membrane and the lower coverslip create a flow cell for bacterial cultivation. When air pressure is applied to deform the membrane downward, the cylindrical protrusions will directly compress cells beneath them. By adjusting the air pressure value, the mechanical force exerted on cells can be flexibly and rapidly altered.

**Figure 3:**
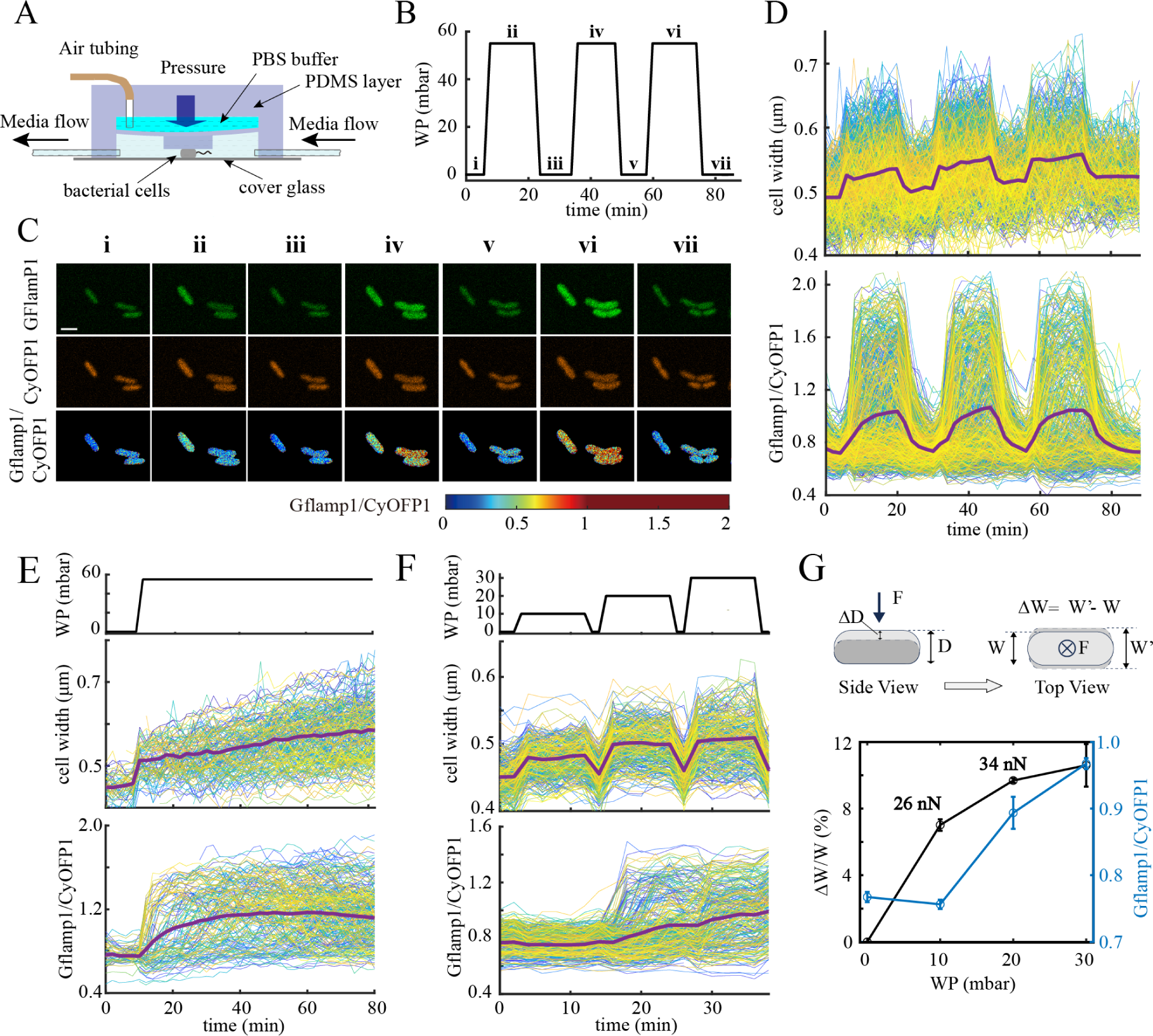
Direct Mechanical Compression at Single-cell Level induces cAMP signaling. (A) Schematic diagram of the microfluidic device designed for direct bacterial compression. (B-D), Experimental results from compressing bacterial cells using a cyclic working pressure curve (B). WP, working pressure. (C) Representative time-lapse images showing Gflamp1, CyOFP1, and Gflamp1/CyOFP1 signals. Scale bar: 2 µm. Roman numerals correspond to time points in (B). (D) Compiled single-cell data (thin lines) and average values (thick magenta line) for changes in bacterial width and Gflamp1/CyOFP1 signal. Different colors of thin lines represent data from different individual cells. (E) Time-series values of bacterial width and Gflamp1/CyOFP1 signal changes when bacteria are subjected to sustained compression force. (F) Time-series values of bacterial width and Gflamp1/CyOFP1 signal changes in bacteria under increasing working pressure. (G) Schematic illustration of bacterial deformation under compression and the calculated force curve based on deformation data from (F). The black curve shows the average relative deformation at different working pressures, estimated compression forces exerted on bacteria were marked near corresponding data points. The blue curve represents the cell-average Gflamp1/CyOFP1 signal at different working pressures.

We designed a cyclic working pressure profile alternating between 55 mbar for 16 minutes and 0 mbar for 10 minutes, repeating for three cycles (Fig. 3B), with fluorescence images captured every two minutes. Microscopic observations revealed that bacteria widened under high pressure (Fig. 3C, Movie S5), confirming successful pressure-induced deformation. Concurrently, the Gflamp1/CyOFP1 signal is increased under high pressure, and both the cell width and Gflamp1/CyOFP1 signal return to baseline levels when the pressure is removed (Fig. 3C). To perform quantitative analysis on multiple bacterial cells, we integrated the Omnipose cell recognition tool with an image tracking algorithm ^28^, allowing us to perform temporal tracking of Gflamp1/CyOFP1 and cell width values and aggregate data from over 300 single-cell sequences. Strikingly, we observed that both the average Gflamp1/CyOFP1 and the cell width curves exhibited perfect synchronization with the working pressure curve (Fig. 3D). Bacterial width responded almost instantaneously to pressure changes, increasing by ∼9% after compression, while the Gflamp1/CyOFP1 signal gradually increased from 0.8 to 1.1 within 6-10 minutes post-compression and decayed exponentially to baseline within 10 minutes after pressure removal (Fig. 3D). The 6-10 minute response time is roughly consistent with the kinetics of cAMP degradation by CpdA ^22^. Repeating the experiments using Gflamp1:R307W abolished the Gflamp1/CyOFP1 response while preserving the compression-induced width changes (Fig. S7). These results demonstrate that using mechanical compression as the sole stimulus is sufficient to elevate bacterial intracellular cAMP levels, ruling out the dependence of pressorum sensing on intercellular transmission of surface molecules or other contact-sensing mechanisms. We next performed the same single-cell level mechanical compression experiments on *cyaA*, *cyaB*, *pilJ*, and *chpA* mutant strains. Consistent with the former data from the agarose chamber experiments, only the *cyaA* strain exhibited a Gflamp1/CyOFP1 signal response to the applied pressure (Fig. S7).

### Bacteria Maintain Elevated Intracellular cAMP Levels Under Persistent Mechanical compression

The single-cell compression device allows for flexible application of various input signal waveforms to investigate the dynamic response of the Pil-Chp system, which is homologous to the flagellar chemotaxis system in *E. coli* ^29^. A key feature of the chemotaxis system is its adaptability, where the output signal gradually decays to baseline levels upon persistent external stimuli ^30^. To explore the adaptability of the Pil-Chp system, we increased the working pressure from zero to 55 mbar and maintained it at 55 mbar. The bacterial width rapidly increased by 10% after initial compression, indicating compression-induced deformation (Fig. 3E). During the subsequent sustained high-pressure phase, the bacteria slowly widened (Fig. 3E), similar to the gradual widening observed in the cyclic high-low pressure experiments (Fig. 3D), probably due to force-induced plastic deformation in cell wall ^31^. Surprisingly, the Gflamp1/CyOFP1 values remained elevated after the initial response, showing no decay for at least 60 minutes, close to the generation time of bacteria, under high pressure (Fig. 3E). This suggests that the Pil-Chp system may lack the adaptability in cAMP synthesis. The adaptability of the *E. coli* flagellar chemotaxis system relies on the methylation and demethylation of MCPs by CheR and CheB ^32^, respectively, corresponding to the actions of PilK and ChpB on PilJ in the Pil-Chp system. The role of methylation in the Pil-Chp system warrant further investigation.

### Activation of cAMP Signaling Requires a Compressive Force on the order of 10 nN

To determine the threshold of mechanical force to trigger the cAMP response, we incrementally increased the working pressure in steps of 10 mbar, maintaining each step for 10 minutes before returning to zero pressure for one minute and then proceeding to the next step (Fig. 3F). Returning to zero pressure between steps allowed for measurement of the cell width before compression. We observed that the Gflamp1/CyOFP1 signal started to increase when working pressure changes from 10 mbar to 20 mbar, corresponding to a 7% to 9% increase in average cell width (Fig. 3F, 3G).

Schematics shows the deformation of bacterial cell in the compression experiments (Fig. 3G). The PDMS pillar compressed bacteria from above to generate an indentation on cell. Concurrently, an increase in cell width was observed in microscopic images, which capture the top view of cells. A bacterial cell is modelled as a prolate ellipsoidal elastic shell with long axis *L*_0_, short axis *W*_0_, and internal turgor pressure *p*. The mechanical force *F* exerted at the top middle point of the long axis of a bacteria can be estimated by ^33^:

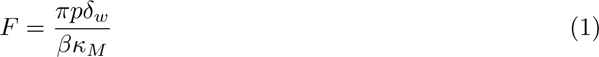

swhere *β* = *δ_w_/δ_D_* is the ratio of cell width variation *δ_w_*to cell height variation *δ_D_*, and *κ_M_* = (*κ_L_* + *κ_W_*)/2 is the mean curvature with 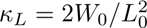 and *κ_W_* = 2/*W*_0_.

For a typical bacteria cell, we assume approximately *L*_0_ = 2 *µ*m, *W*_0_ = *D*_0_ = 0.5 *µ*m, *p* = 0.5 MPa ^34^ and *β* = 1 (see Supplementary Information). The estimated mechanical force for 7% to 9% change in cell width was 26∼34 *nN*, two orders of magnitude greater than the pulling force of type IV pili (100 pN) ^35^. This result is consistent with the predicted compressive force required to activate cAMP signaling of cells grown in the agarose gel chamber (Fig. 1G).

### Formation of Traveling Ring Patterns of Elevated Intracellular cAMP in Compressed Bacterial Colonies

Autoinducers in quorum sensing systems are self-generated molecules that diffuse into surroundings. When combined with appropriate spatial boundary conditions and genetic feedback loops, bacteria can exhibit coordinated gene expression behavior on a large scale, forming intriguing spatial patterns ^36–38^. However, the mechanical force signals in pressorum sensing have distinct generation and release characteristics compared to autoinducers. This prompted our curiosity about the spatiotemporal response of pressorum sensing on a macroscopic scale.

We allowed a single bacterial cell to grow axisymmetrically into circular colonies on a glass slide covered by a thick agarose hydrogel, monitoring the evolving spatial distribution of cellular cAMP levels in the colony. Compressed by the hydrogel blanket, the colony grew epitaxially and inserted into glass-hydrogel interface. The spatial distribution of colony thickness revealed the formation of multiple layers with steep transitions between neighboring layers, creating step-like features at the colony-hydrogel interface (Fig. 4A). A striking phenomenon was the spatial overlapping and synchronous traveling of Gflamp1/CyOFP1 ring patterns and steps (Fig.4A, 4B), with a consistent spacing of ∼ 23 µm between the two steps/rings (Fig. 4B). Confocal microscopy cross-section images further confirmed the colony’s step-wise structure (Fig.4C, Movie S6), revealing discontinuous incrassation in thickness that form cell plateaus separated by steps.

**Figure 4:**
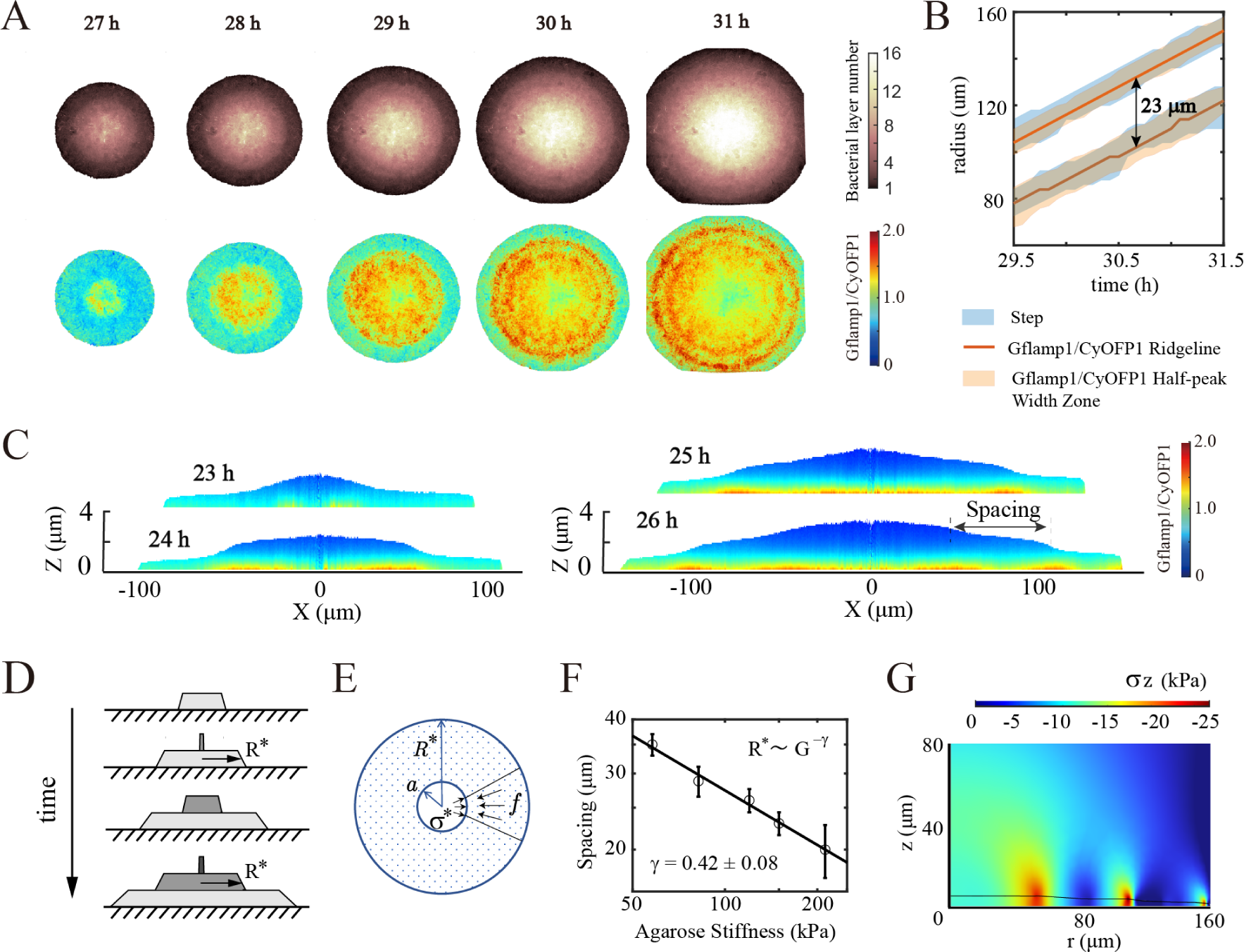
Formation of traveling ring patterns of elevated intracellular cAMP in compressed bacterial colonies. (A) Temporal evolution of colony thickness and spatial distribution of Gflamp1/CyOFP1 signal at the interface between 3.0% agarose gel and glass. (B) Time-dependent curves of the radial position of colony steps (blue shaded area) and the peak position of Gflamp1/CyOFP1 radial distribution (orange solid line). The orange shaded area indicates the radial region where the Gflamp1/CyOFP1 signal intensity exceeds half of its maximum value. The radial distance between two Gflamp1/CyOFP1 peak positions is indicated by a line with double-headed arrows. (C) The *z*-direction scan images of colony cross-sections at different time points for a bacterial colony growing at the interface between 2.0% agarose gel and glass. (D) Schematic model of step formation during colony growth. (E) Force diagram illustrating the upward compression of a laterally expanding bacterial colony leading to new layer formation. *R^∗^* and *σ^∗^* denote the critical colony radius and internal pressure at new layer formation, respectively. *f* represents the frictional force opposing outward colony expansion, and *a* indicates the critical yield size in the central region of the colony. (F) Relationship between gel stiffness and step spacing plotted on a double logarithmic scale, with the black solid line representing linear fitting results. (G) Numerical simulation results of compressive stress in the *z*-direction experienced by step-growing colonies under gel compression.

We proposed a simple model to explain the step-wise growth of the circular bacterial colony, as illustrated in the schematic diagram (Fig. 4D). In the early stage, the first colony layer expanded axisymmetrically and was subjected to centripetal resistant stress *τ* along the colony-glass and colony-hydrogel interfaces. The resulted radial compression force diverged towards the center and scaled with the radius of colony ^39^. At the critical radial compression stress *σ^∗^* and radius *R^∗^*, the cells near the center yielded and were pushed vertically to initiate the formation of the second layer ^40^. The consistent spacing between the two steps indicates the equal radial expansion speed of the two layers (Fig. 4B), then the first and the second layer reach the critical radii of 2*R^∗^* and *R^∗^* simultaneously, and the cells near the center of the second layer again yield and were pushed vertically to form the third layer. Thus the spacing between two steps is *R^∗^*. Scaling analysis shows that *R^∗^* = *a*(*σ^∗^/G*)*^γ^*, where *G* is the shear modulus of agarose hydrogel, *a* is the radius of yielding cells near the center (Fig. 4E). Using hydrogels of different concentrations, we saw the scaling exponent *γ* = −0.42 *±* 0.08 (Fig. 4F). For the first layer with frontal step of height *h* and width *δ* (Fig.4D), the vertical compression stress *σ_⊥_* on colony is linear with shear strain *h/δ*, and further proportional to the centripetal resistant stress *τ*, i.e. *σ_⊥_ ∼ Gh/δ ∼ τ*. At the critical point, the balance between the radial force of the central yielding cells and the centripetal resistant force requires *σ^∗^ah* = *τ* (*R^∗^*^2^ −*a*^2^)/2, such that *R^∗^ ∼ G^−^*^1^*^/^*^2^. The scaling exponent *γ* = −0.5 is in well agreement with the experimental value.

While the dynamic process giving rise to the traveling Gflamp1/CyOFP1 ring patterns remains to be clarified, our findings suggest a strong correlation between stress concentration at steps and the formation of ring patterns. As the growing colony slid over the glass surface, bacteria that attached firmly to the glass surface may sustain frictional forces continuously, explaning the observed signifciant Gflamp1/CyOFP1 signals at the colony-glass interface (Fig. 4G). Notably, the peaked friction forces may be triggered by the concentrated stresses under steps, where the strongest Gflamp1/CyOFP1 signals were seen moving with steps (Fig. 4A,C). To complement this correlation, a *pelpsl* mutant strain with impaired extracellular polysaccharide production and consequently reduced cell-cell and cell-glass interactions showed neither colony steps nor Gflamp1/CyOFP1 ring patterns (Fig. S8, Movie S6, S7).

## Discussion

In this study, we found that when *P. aeruginosa* grows in a confined space and reaches a densely-packed state, compressive forces between bacteria trigger a cAMP response, upregulating T3SS expression. We introduced the term ‘pressorum sensing’ to describe this phenomenon, which is analogous to ‘quorum sensing.’ Both processes involve a phenotypic switch of the entire bacterial population, triggered by the accumulation of a signal as the population density increases. However, quorum sensing and pressorum sensing differ in their signal types and specificity. Quorum sensing uses self-generated, species-specific chemical autoinducers, allowing the distinction between self and non-self populations. In contrast, pressorum sensing is triggered by mechanical compression and cannot differentiate the origin of the growth-induced compression.

Pressorum sensing provides a potential mechanism for *P. aeruginosa* to activate the T3SS within a host. This mechanism ensures that bacteria transition to a virulent state only when their population is sufficiently large and growing normally, reducing the risk of premature detection and elimination by the immune system. Additionally, pressorum sensing may help bacteria recognize spatially restricted colonization environments, such as tissue interstices, allowing appropriate activation of the T3SS. Another signal that activates T3SS expression within a host is low calcium concentration, which regulates ExsA translation through the LadS/GacS-GacA-RsmYZ-RsmA cascade ^41,42^. It is possible that pressorum sensing acts synergistically with the low calcium signal, particularly in environments such as macrophage phagosomes, leading to a significant enhancement of T3SS expression.

Pressorum sensing is also a type of bacterial mechanosensing that requires mechanical forces of ∼10 nN. Other mechanosensing mechanisms in *P. aeruginosa* include: (1) Rheosensing, where bacteria sense the surrounding flow field rather than shear force to regulate gene expression ^43^; (2) PilY1-mediated surface sensing, where the mechanical sensing protein PilY1 at the tip of type IV pili transmits signals to modulate FimS-PilJ and PilO-SadC interactions, altering intracellular c-di-GMP concentration and minor pilin expression ^44–46^; and (3) Twitching motility-mediated surface sensing, where type IV pili become loaded upon surface adhesion, altering PilA-PilJ interaction through extension and retraction, regulating PaQa gene operon expression ^11,12^. This mechanism allows bacteria to perceive surface attachment and stiffness. Twitching motility-mediated surface sensing and pressorum sensing share similarities in their upregulation of intracellular cAMP levels through the Pil-Chp system. Howerver, they differ in their reliance on type IV pili mechanics and the magnitude of forces involved. pressorum sensing does not require pili extension and retraction and involves forces two orders of magnitude higher than the pilus retraction force (100 pN).

From a synthetic biology perspective, pressorum sensing provides a unique approach to translating mechanical force signals into bacterial phenotypic responses. Two promising routes for additional investigation are presented by this finding. First, by controlling the physical boundaries of bacterial growth, for example through 3D printing techniques, researchers can manipulate the spatial patterns of bacterial gene expression on a large scale or alter the local phenotype by applying mechanical force at specific locations. Second, bacteria themselves may serve as active materials for sensing mechanical forces, offering information about static stress or even stress history. The response dynamics of cAMP to mechanical force can be fine-tuned by adjusting the expression level and activities of the Pil-Chp system and CpdA, while downstream gene expression response to cAMP wave profiles can be modified using designed genetic regulatory circuits. These possibilities highlight the potential for pressorum sensing to open up new avenues in mechanogenetics. As our understanding of pressorum sensing deepens, it may lead to the development of novel synthetic biology tools and applications that leverage the interplay between mechanical forces and bacterial physiology.

## Methods

### Bacterial strains and growth conditions

Bacterial strains and plasmids were listed in Table S1 and S2 in the Supplementary Information. All the strains used in the microscopic experiments underwent a two-step culture process at 37 *^◦^*C. Initially, bacteria were scraped from agar plates and shook overnight, followed by a 50× dilution and subsequent shaking for 6 hours (OD600∼0.5). The culture medium employed was FAB medium supplemented with 30 mM sodium glutamate and 30 µg mL*^−^*^1^ gentamycin (FABG).

### Construction of gene knockout and transcriptional reporter strains

Knockout of target genes in *P. aeruginosa* were conducted using a CRISPR-based method ^47^. Briefly, a pCasPA plasmid was first electroporated into original *P. aeruginosa* strains and the target colonies were selected on LB agar plates supplemented with 50 *µ*g/mL tetracycline. The induction of Cas9 Expression in the resultant strains was achieved via addition of 2% arabinose in LB medium and shook for 4 hours at 37 *^◦^*C before next electroporation step ^48^. Then a pACRISPR plasmid, containing the up-down recombinant fragments and gRNA for target genes, were electroporated into the strains. Colonies were selected on LB agar plates supplemented with 100 *µ*g/mL tetracycline and 150 *µ*g/mL carbenicillin. Successful gene editing was confirmed through PCR analysis. Additionally, the pCasPA and pACRISPR plasmids within the resultant strains were subsequently eliminated via bacterial streaking and selection on no-salt LB agar plates supplemented with 15% sucrose.

To construct the cAMP reporter strain, we assembled the *gflamp1* gene fragment (without the stop codon), with the constitutive promoter *J23100*, the ribosome binding site RBSII, and the *cyOFP1* gene encoding segment, replacing the arabinose-inducible expression gene module araC-pBAD in pJN105, resulting in the gflamp1-pJN plasmid. The gene expression structure is designated as J23100-RBSII-gflamp1-cyOFP1. Subsequently, the plasmid was separately introduced into wildtype PAO1 or various gene knockout mutant strains.

In the construction of the transcriptional reporter plasmids, we initially modified the original pJN105 plasmid by removing the arabinose-inducible gene module araC-PBAD and replacing it with the gene expression module RBSII-sfGFP-T0T1-J23102-CyOFP1, yielding the pJNTR plasmid. RBSII is a strong ribosome binding site, and T0T1 represents a transcription terminator. J23102 (http://parts.igem.org/Part:BBa_J23102) serves as a constitutive promoter controlling the expression of CyOFP1 as an internal standard. All transcriptional reporters were constructed by inserting 500-1000 *bp* upstream sequences of the target gene into pJNTR, right before RBSII, forming the expression structure Px-RBSII-sfGFP-T0T1-J23102-cyOFP1. Then the plasmids were electroporated into *pilQ* or *pilQcyaB* mutant strains. The transcriptional activity of the target promoters was evaluated via the fluorescence ratio of SfGFP to CyOFP1.

### Bacterial incubation in confined agarose chamber

The microchip platforms were fabricated with polydimethylsiloxane (PDMS) (Sylgard 184; Dow Corning) using standard soft lithography methods. Wafers were coated with SU-8 2010 photoresist (MicroChem Inc., Newton, MA, USA) to form film depositions. Silicon wafer was patterned with an array of cylindrical cavities (diameter 30 µm or 60 µm, height 2 µm). A mixture of prepolymer PDMS and cross-linking agent (mass ratio, 10:1), was poured into the molds with the aforementioned silicon wafer as the base, and baked at 80 *^◦^*C for one hour. Subsequently, a small piece of PDMS block (14 mm in length, 10 mm in width) was excised from the mold, with cylindrical protrusions on its underside. The PDMS block was then flipped and placed flat on a surface, with a rectangular pressure block (4 mm in thickness) positioned above, featuring a central through-hole (11 mm in length, 7 mm in width). Ensured that the PDMS block precisely sealed the rectangular pressure block’s through-hole. Subsequently, 400 µL of hot solution of FABG 2.5% agarose gel was poured into the through-hole, and allowed to cool and solidify for 10 minutes. After removal from the mold, the solidified gel blocks were flipped to expose the circular cavities at their base, onto which 1.5 µL of bacterial suspension diluted 3000-fold was added. After air-drying, the agarose pads were flipped and gently pressed onto glass-bottomed culture dishes, sealed, and observed on a microscope.

### Single-cell compression experiments

Fabrication process of wafer molds and microfluidic chips were included in the Supplementary Information. Microfluidic Chips were soaked in deionized water at 4 *^◦^*C for at least two days. Before experiments, connecting the gas and liquid lines. Injecting PBS buffer into the gas line, displacing all air, and then blocking the tailpipe. Injecting 10× diluted bacterial solution into the liquid line, waiting for bacterial adhesion under a microscope, and simultaneously initiating the infusion of FABG medium at a flow rate of 200 µL/min. Once a sufficient number of bacteria have adhered, connecting the gas line to the air pump (brand, model: FluidicLab, PC-1), gradually increasing the pressure to just press the PDMS square compression pillars against the surface of the glass slide (zero working pressure position), and culturing at 30 *^◦^*C for one hour. Then, applying different pressure working curves to compress the bacteria while collecting fluorescence images simultaneously.

### Monitoring intracellular cAMP level in bacterial colonies

The FABG culture medium was supplemented with various concentrations (1.5%∼ 3.0%) of agarose (BBI, low EEO), boiled, poured into culture dishes, and allowed to cool for later use. The harvested bacterial culture of the cAMP reporter strain (OD∼0.5) was diluted 50,000-fold, and then two micro-liters were dispensed onto agarose gel patches (1.0 cm x 1.0 cm). After air-drying, the patches were gently pressed onto glass-bottom dishes and sealed. The prepared samples were then incubated at room temperature. Subsequently, fields containing bacterial colonies were selected under a micro-scope, and fluorescence images of Gflamp1 and CyOFP1 were captured using a 40× objective every 5 minutes.

### Fluorescent image acquisition and data analysis

The agarose chamber experiments (including observations of Gflamp1 and transcriptional reporters) and large colony experiments were conducted using an Olympus IX83 inverted wide-field fluorescence microscope. Fluorescence imaging was achieved through excitation with a 488 nm laser, with simultaneous acquisition of SfGFP/Gflamp1 and CyOFP1 images using a dual-camera setup (Photometrics BSI). Corresponding filter sets were FF01-515/30-25 (Semrock) and FF01-598/25-25 (Semrock). The agarose chamber experiments utilized a 100× objective, while the large colony experiments utilized a 40× objective. Fluorescence images were captured every five minutes. Image processing was conducted using self-written MATLAB code as previously reported ^49^. Briefly, SfGFP/Gflamp1 and CyOFP1 images captured by two cameras were aligned using a built-in function (imwarp), and crosstalk of fluorescence were corrected using a cross-talk matrix. Cell mask images were determined from CyOFP1 images using a fixed threshold, and the corresponding SfGFP/Gflamp1 and CyOFP intensities were calculated using a built-in function (regionprops). Z-stack images of colonies were captured using a spinning-disk confocal microscope (Nikon Microscope, YOKOGAWA CSU-W1 spindisk). Each stack consisted of 101 images taken at 0.1 µm intervals. Scans were performed every 30 minutes.

Single-cell compression experiments employed a spinning disk confocal microscope (Leica MM165), equipped with a 100× objective and dual cameras (Photometrics BSI) for simultaneous acquisition of Gflamp1 and CyOFP1 images. Filter sets used were B525/50 and B617/73 (Semrock). Fluorescence images were captured every two minutes. The image processing method consists of three parts: (1) alignment and color correction of dual-camera images, following the method described above; (2) single-cell identification using CyOFP1 images, employing the omnipose tool based on the Anaconda environment, with the ‘bact fluor omni’ model selected and other parameters set to default values; (3) single-cell tracking performed using a MATLAB algorithm developed in-house, using overlapping regions in adjacent frames to determine the same bacterial cell.

## Supplementary information

Additional results, additional methods, additional tables and figures can be found in Supplementary Information.

## Acknowledgements

We extend our gratitude to Professor Jun Chu for generously providing the cAMP reporter protein Gflamp1. We thank JiZhou Huang for introducing the Omnipose tool for image processing, Jinjuan Wang for her help in microfluidic device fabrication, and Jialun Lin for his assistance in formatting the manuscript. This work was supported by National Key Research and Development Program of China (Grant No. 2020YFA0906900 to Fan Jin), the National Natural Science Foundation of China (Grant No. 32101177 to Yajia Huang), National Key Research and Development Program of China (Grant No. 2023YFB4605403 to Haiyi Liang), Shenzhen Engineering Research Center of Therapeutic Synthetic Microbes (Grant No. XMHT20220104015 to Fan Jin).

## Author Contributions

We were responsible for this study as follows: conceptualization, Fan Jin; Primer design for strain construction, Yajia Huang; Strain construction, Jiarui Xiong and Wenwen Xiao; Initial experimental exploration, Yajia Huang and Jiarui Xiong; Postulation of the mechanical compression dependent cAMP signaling mechanism, Fan Jin; Experimental design, Fan Jin, Lei Ni, Yajia Huang; Stainig of type IV pili, Yajia Huang; Acquisition of data for bacteria grow in microcavities, Yajia Huang and Lei Ni; Microfluidic device design, Yue Yu; Microfluidic device fabrication, Lei Ni and Hui Wen; Direct bacterial compression experiments, Lei Ni and Yajia Huang; Colony experiments, Yajia Huang; Image processing and data presentation, Lei Ni; Theoretical explanation of colony step formation and cAMP traveling ring pattern at steps, Haiyi Liang and Fan Jin; Simulation and calculation of forces on bacteria in agarose hydrogel and during compression, Haiyi Liang and Yaoxin Huang; Manuscript drafting: Haiyi Liang (traveling ring pattern formation in colony); Yaoxin Huang (physical modeling of forces acting on bacteria); Lei Ni (remaining content); Manuscript revision and review, Fan Jin, Lei Ni, Haiyi Liang, Yajia Huang and Yaoxin Huang.

## Declarations

The authors declare no competing interests.

