## Supplementary Information for "Pressorum Sensing: Growth-induced Compression Activates cAMP Signaling in *Pseudomonas aeruginosa*"

### Contents

**Additional Results and Methods** **3–11**

**Additional References** **12**

**Supplementary Figures S1-S9** **13–21**

**Supplementary Movies S1-S7** **22–28**

**Table S1-S2** **29–38**

### Additional Results and Methods

#### Initial observations of Gflamp1/CyOFP1 signals of *P. aeruginosa* cells grown between hydrolgel-glass interface

We started with an experimental setup in which wild type *P. aeruginosa* cells were allowed to grow between the crevice of an agarose gel and a coverglass. Initially, the bacterial cells exhibited twitching motility across the interface, with substantial cell-to-cell heterogeneity in the Gflamp1/CyOFP1 signal , wherein fluctuations could be observed within certain individual cells. We noted an elevated Gflamp1/CyOFP1 signal at the expanding periphery of the twitching colony, while lower levels were observed at the center (Fig. S1A). At a later stage, the bacteria transitioned into rapid, fluidic-like motility across the entire interface, accompanied by an overall attenuation of the Gflamp1/CyOFP1 signal (Fig. S1B, movie S1 (wild type 1.5%)). Remarkably, as the cells gradually became densely packed and traffic-jammed, we observed a global elevation in the Gflamp1/CyOFP1 (Fig. S1A, S1B, and movie S1). In experiments using higher concentrations of agarose gel and *chpB* or *fimS* knockout strains, we observed similar increases in Gflamp1/CyOFP1 after bacterial traffic jams (Fig. S1C). However, this phenomenon disappeared in the *cyaB* knockout strain (Fig. S1C). In *pilA*, *pilB*, and *pilT* knockout strains, which lack twitching motility, the fluidic-like motion stage was absent, along with the absent of global decrease of Gflamp1/CyOFP1 after transitioning into the fluidic-like stage (movie S1). Instead, bacteria formed gradually growing colonies that compressed and mixed with each other, suggesting that type IV pili mediate the fluidic-like motion. Notably, during the mixing and expansion of these colonies, we observed coordinated changes in intracellular Gflamp1/CyOFP1 levels across large spatial scales. (Fig. S1D, movie S1).

#### Experimental procedure for initial observation of *P. aeruginosa* cells cultured at the hydrogel-glass interface

The *P. aeruginosa* cells was cultured overnight in 1 mL of FABG medium at 37 °C. The culture was then diluted 50× in fresh FABG medium and grown for an additional

6 hours at 37 °C until reaching an OD of approximately 0.5. The bacterial suspension was further diluted 50-fold in fresh FABG medium. A 2 µL aliquot of this dilution was deposited onto a 1.5% agarose gel pad which was made using FABG medium. After air-drying, the gel pad was inverted and placed in a round-bottom culture dish with a cover slip base. Observations were conducted using an Olympus IX83 inverted fluorescence microscope, equipped with the same lens configuration as described in the methods section of main text. Images were captured at 1-minute intervals. During microscopic observation, samples were maintained at 30 °C.

#### Experimental procedure for co-culturing *P. aeruginosa* and *E. coli* cells at the hydrogel-glass interface

*E. coli* and *P. aeruginosa* cells were individually cultured overnight in 1 mL of M9CA medium (Sangon Biotech) supplemented with 30 µg mL$^{-1}$ gentamicin (M9CAG) at 37 °C. Each culture was then diluted 100-fold in fresh M9CAG medium and grown to an OD of approximately 0.5 at 37 °C. The bacterial suspensions were harvested and mixed at a ratio of 10:1 (*E. coli* : *P. aeruginosa*). A 2 µL aliquot of this mixture was deposited onto a 1.5% agarose gel pad prepared with M9CAG medium. After air-drying, the gel pad was inverted and placed in a round-bottom culture dish with a cover slip base. Observations under microscope are the same as described for initial observation of *P. aeruginosa* cells cultured under agarose hydrogel.

#### Staining of Type IV pili in *pilQ*, *pilT* and wild type *P. aeruginosa* strains

Bacterial cultures were grown overnight in FABG medium and subsequently diluted 50-fold in fresh FABG medium for 2 hours before supplemented with arabinose. The cultures were then shaked for an additional 4 hours at 37 °C before harvested. 200 µL of bacterial suspension was taken and directly mixed with 2 µL of Alexa Fluor$^{TM}$ 488-maleimide (Thermo A10254) dye solution (10 mg mL$^{-1}$ in DMSO). The mixture was gently shaked at 400 rpm and incubated at 37 °C in a metal bath for 15 minutes. Then, bacterial cells were washed three times with fresh FABG medium (3000 rpm for 5 minutes each time). Bacterial pellet was resuspended in 1 mL of FABG medium and 1.5 µL of the suspension was dropped on a 0.5% agarose gel pad made with FABG medium. The pad was turned upside down on a cover glass-bottom dish for observation under microscope.

**Fabrication of microfluidic chips for direct compression of bacterial cells**

The air-layer wafer was coated with SU-8 2010 photoresist to form film deposition of 20 µm thickness. The liquid-layer wafer consists of two layers: the first layer employs SU-8 6002 photoresist to create a thin film with a thickness of approximately 3.5 µm, corresponding to the red square region in Fig. S6. This layer forms 3.5 µm square protrusions on the template. The second layer utilizes SU-8 2010 photoresist to form a film with a thickness of 18 µm, corresponding to the black line region in Fig. S6, where the square compression pillars and circular support pillars are recessed areas. Consequently, the final height of the pressure pillars is 14.5 µm, while the height of the support pillars is 18 µm. In this way, 3.5 µm gaps were created between the compression pillars and the cover glass at the bottom of the chip, thus preventing adhesion during bonding with cover glass. The photolithography procedures were conducted following the manufacturer's guidelines for respective photoresists.

Sebsequent process of microfluidic chip fabrication involves: (1) Mixing PDMS pre-polymer with a cross-linking agent in a 10:1 ratio (20 g+2 g), pouring it into air-layer mold, curing at 80 °C for one hour, followed by cutting into squares and punching holes to obtain the upper layer chip. (2) Mixing PDMS pre-polymer with a cross-linking agent in a 9:1 ratio(9 g+1 g), pouring it onto the liquid-layer wafer mold, spinning at 800 rpm for 30 seconds, baking at 80 °C in a flat metal heating platform for one hour to obtain the PDMS membrane. (3) Plasma cleaning the upper layer chips and PDMS membrane (together with the liquid-layer wafer mold) for 50 seconds, then add 100 µL of methanol onto the PDMS membrane, quickly adhering the upper layer chip under an optical microscope, make sure the square pillars of the PDMS membrane are aligned with the central region of the square chambers in the upper layer chip. After allowing methanol to evaporate for 15 minutes, vacuuming for 3 minutes, and baking at 80 °C for two hours. Peeling off the entire chip from the liquid-layer wafer mold, punching holes, bonding the bottom layer of the chip (outer side of the membrane) to a cover glass ($24\,\text{mm} \times 50\,\text{mm}, 0.17\,\text{mm}$ thick) (plasma cleaning for 50 seconds, baking at 80 °C for one hour).

##### Sequence of CyOFP1-Gflamp1 fusion protein

The underlined characters represent the amino acid sequences of CyOFP1, bold characters indicate the amino acid sequences of Gflamp1, and italic characters denote the fusion protein linker sequences.

MVSKGEELIKENMRSKLYLEGSVNGHQFKCTHEGEGKPYEGKQTNRIKVVEGGPLPFAFDILATHFMYGSKVFIKYPADLPDYFKQSFPEGFTWERVMVFEDGGVLTATQDTSLQDGELIYNVKVRGVNFPANGPVMQKKTLGWEPSTETMYPADGGLEGRCDKALKLVGGGHLHVNFKTTYKSKKPVKMPGVHYVDRRLERIKEADNETYVEQYEHAVARYSNLGGGMDELYK*GGGGMGSHHHHHHGMASMTGGQQMGRDLYDDDDKDP***MGFYQEVRRGDFVRNWQLVAAVPLFQKLGPAVLVEIVRALRARTVPAGAVICRIGEPGDRMFFVVEGSVSVATNWGNVYITADKQKNGIKANFKIRHNVEGGGVQLAYHYQQNTPIGDGPVLLPDNHYLSVQSKLSKDPNEKRDHMVLLEFVTAAGITLGMDELYKGGTGGSMVSKGEELFTGVVPILVELDGDVNGHKFSVRGEGEGDATNGKLTLKFICTTGKLPVPWPTLVTTLTYGVQCFARYPDHMKQHDFFKSAMPEGYIQERTIVFKDDGTYKTRAEVKFEGDTLVNRIELKGIDFKEDGNILGHKLEYNRVNPVELGPGAFFGEMALISGEPRVATVSAATTVSLLSLHSADFQMLCSSSPEIAEIFRKTALERRGAAASA**

#### Modeling of mechanical stress on bacteria under gel

| | | | |
|---|---|---|---|
| Reference state coordinate | $\mathbf{X}$ | Nominal stress | $s$ |
| Current state coordinate | $\mathbf{x}$ | True stress | $\sigma$ |
| Deformation gradient | $\mathbf{F}$ | Number of polymer chains | $N$ |
| Current state velocity | $\mathbf{v}$ | Boltzmann constant | $k$ |
| Body force | $\mathbf{b}$ | Temperature | $T$ |
| Surface force | $\mathbf{t}$ | Flory–Huggins parameter | $\chi$ |
| Energy functional | $\Pi$ | Polymer mass fraction | $\psi$ |
| Helmholtz free energy | $W$ | Molecule volume | $\Omega$ |
| Chemical potential | $\mu$ | Inverse of $\mathbf{F}$ | $\mathbf{H}$ |
| Solvent nominal concentration | C | Inverse of $\mathbf{R}^s$ | $\mathbf{M}$ |
| Solvent flux | $I$,$\mathbf{J}$ | Growth coefficient | $k_g$ |
| Unit outward normal | $\mathbf{N}$ | Friction coefficient | $k_f$ |
| Dissipation tensors | $\mathbf{R}^v$, $\mathbf{R}^s$ | Spring coefficient | $k_s$ |

##### Polymeric gels model

The polymeric gel model established in Hong and Wang is used in our simulation[1,2]. We only look at simple gels with only water molecules as the mobile solvent molecules. The dry network under no mechanical load is taken as the reference state, and each marker is named using its coordinate $\mathbf{X}$ in the reference state. At time $t$, marker $\mathbf{X}$ moves together with the host polymer particle to a location with coordinates $\mathbf{x}(\mathbf{X}, t)$. The deformation gradient is $\mathbf{F} = \frac{\partial \mathbf{x}}{\partial \mathbf{X}}$.

Assuming that the solvent nominal concentration $C$ is related to the volumetric expansion det$\mathbf{F}$ and that the energy is dissipated only through solvent migration and viscous deformation, the energy functional with a constraint term for the conservation of the number of solvent molecules via the Lagrange multiplier $\mu(\mathbf{X}, t)$ is written as

$$
\begin{aligned}
\hat{\Pi} =& \int_V \tfrac{\partial W}{\partial t}\,\mathrm{d}V - \int_V b_i v_i\,\mathrm{d}V - \int_{\partial V} t_i v_i\,\mathrm{d}A - \int_{\partial V} \bar{\mu} I\,\mathrm{d}A \\
&+\tfrac{1}{2}\int_V R^v_{iKjL}\tfrac{\partial v_i}{\partial X_K}\tfrac{\partial v_j}{\partial X_L}\,\mathrm{d}V \\
&+\tfrac{1}{2}\int_V R^s_{KL} J_K J_L\,\mathrm{d}V \\
&+\int_V \mu\left(\tfrac{\partial C}{\partial t} + \tfrac{\partial J_L}{\partial X_L}\right)\mathrm{d}V,
\end{aligned}
\tag{1}
$$

where $W$ is the Helmholtz free energy density determined by $\mathbf{F}$ and $C$, $\mathbf{v} = \frac{\partial \mathbf{x}}{\partial t}$ is the velocity, $\mathbf{b}$ and $\mathbf{t}$ respectively are the body force acting throughout the bulk and the surface force acting on the boundary, $\mathbf{R}^v$ and $\mathbf{R}^s$ are the generalized viscosity dissipation tensors due to viscous deformation and solvent migration respectively, $\mathbf{J}$ is the diffusion flux of solvent and $I = \mathbf{J} \cdot \mathbf{N}$ on the surface where $\mathbf{N}$ is the unit outward normal of the boundary in the reference state. Minimizing the functional $\hat{\Pi}$, the governing equations of the system and the boundary conditions are obtained:

$$\frac{\partial s_{iK}}{\partial X_K} + b_i = 0, \tag{2}$$

$$\frac{\partial C}{\partial t} + \frac{\partial J_L}{\partial X_L} = 0, \tag{3}$$

$$J_K = -M_{KL}\frac{\partial \mu}{\partial X_L}, (\mathbf{M} = (\mathbf{R}^s)^{-1}), \tag{4}$$

in the bulk and

$$s_{iK} N_K = t_i, \tag{5}$$

$$\mu = \bar{\mu}, \tag{6}$$

on the surface, where

$$s_{iK} = \frac{\partial W}{\partial F_{iK}} + R^v_{iKjL}\frac{\partial v_j}{\partial X_L} - \frac{\mu}{\Omega} H_{Ki} \det \mathbf{F}, (\mathbf{H} = \mathbf{F}^{-1}) \tag{7}$$

is the nominal stress, and $\Omega$ is the volume occupied by each molecule in pure solvent. The form of the Helmholtz free energy of the hydrogel is given by

$$W(\mathbf{F}, C) = \frac{1}{2}NkT(F_{iK}F_{iK} - 3 - 2\log(\det\mathbf{F})) - \frac{kT}{\Omega}\left[\Omega C\log\left(1 + \frac{1}{\Omega C}\right) + \frac{\chi}{1+\Omega C}\right], \quad (8)$$

where $N$ is the number of polymer chains per unit volume in the reference state, $kT$ is the temperature in the unit of energy and $N$ is the Flory–Huggins parameter for the enthalpy of mixing. The equations above constitute an initial-boundary-value problem for the unknown field $\mathbf{x}(\mathbf{X}, t)$ and $\mu(\mathbf{X}, t)$.

#### Numerical implementation

In all numerical simulations, a nondimensionalization was taken through normalizing all stress by $\frac{kT}{\Omega} = 4 \times 10^7$ Pa at room temperature and all length by $L_0$ =1 µm. $\mathbf{R}^v$ and $\mathbf{R}^s$ are reduced to isotropic for simplicity and these dynamic parameters have no effect on the following stationary results. 2D axisymmetric components was adopted in all of the simulations.

The weak form of the governing equations with the dimensionless expression for nominal stress is implemented in finite-element method. In all calculations, implicit time discretization and a direct linear solver are used. The spatial interpolation of the displacement field is set as one order higher than that of the chemical potential for matching the element order between stress and chemical potential.

#### Uniaxial compression and stretch of cylindrical hydrogel

We performed unconstrained uniaxial compression and stretch simulations to check the apparent modulus of the hydrogels. The initial geometry and boundary conditions being used in the calculation are shown in Fig. S9A. We set no body force and the hydrogel initially swollen and has reached equilibrium with the surrounding solvent, that is, $\mathbf{b} = 0$ and the initial chemical potential of the gel $\mu_0 = 0$. The final height of the cylinder at a large enough $t$ was set as $h_{\infty,\mathrm{com}}$ =0.1 µm in compression and $h_{\infty,\mathrm{str}}$ =5 µm in stretch. The stress-strain curve in Fig. S9B were obtained for polymer mass fraction

$\psi = 2.5\%$ and the Flory–Huggins parameter $\chi = 0.15$. The elastic modulus at the origin is 378 kPa, in agreement with the experimental data in Normand[3].

**Simulation of force acting on bacterial cells filling microcavities**

The initial geometry with boundary conditions used in the calculation are shown in Fig. S9C. The boundary between the bacteria and the hydrogel is only the boundary of the diffusion equation Eq. 3, not the boundary of the force balance equation Eq. 2. The hydrogels tested in above subsection **'Uniaxial compression and stretch of cylindrical hydrogel'** were used, and a linear elastic material with a Young's modulus of 300 kPa and a Poisson's ratio of 0.3 was used for the bacteria.

We applied growth on the bacteria throughan inelastic deformation gradient with $(1, 1, \gamma_z)$ in the principal directions. The rate of growth was set as $\dot{\gamma}_z = k_g\left(\sigma_z - \sigma_z^*\right)\frac{kT}{\Omega}$ and we chose the growth coefficient $k_g = 1$ where $\sigma_z$ is the $z$-component of true stress and $\sigma_z^*$ is an equilibrium stress[4]. It means that the $z$-directional stress within the bacteria will eventually stabilize uniformly at $\sigma_z^*$. Fig. S9D shows the relationship between $\sigma_z^*$ and the deformation magnitude of the microcavity.

**Simulation of stress concentration at steps of bacterial colonies**

The initial geometry with boundary conditions used in the calculation are shown in Fig. S9E. Contact pairs were set up at the contact boundaries between the bacteria and the hydrogel, where the boundaries of bacteria are set as source boundaries and ones of hydrogel are set as destination boundaries. The materials and initial states of the hydrogel and bacteria are the same as in above subsection **'Simulation of force acting on bacterial cells filling microcavities'**. Radial force $t_r = k_f\sigma_z$ was applied at the bottom of the bacteria as friction only when $\sigma_z < 0$ and we set the friction coefficient $k_f = 0.05$. Simple springs with $k_s = 50$ kPa were applied to the bottom of the hydrogel for adhesion between the hydrogel and the substrate. A homogeneous growth in $z$ direction from 0.25 µm to a 5 µm was applied to the bacteria and another growth in the form above subsection **'Simulation of force acting on bacterial cells filling microcavities'** with $k_g = 10$ and $\sigma_z^* = -30$ kPa was applied only when

$\sigma_z < -30\,\text{kPa}$. The $z$-component of true stress field in the equilibrium state of the system and the $r$- and $z$-components of true stress in the bottom layer of the bacteria are shown in Fig. S9F and Fig. S9G, respectively.

#### Derivation of $\beta$ in equation (1) of the main text

We assume that the elastic shell remains ellipsoidal during deformation and that the surface area and the longest semi-axis remain constants. Let $a_0, b_0, c_0 (a_0 \geq b_0 \geq c_0)$ be initial principal semi-axes of the ellipsoid and deformed to $a, b, c$ . The surface area can be calculated by

$$S(a,b,c) = 2\pi c^2 + \frac{2\pi ab}{\sin(\varphi)} \left( E(\varphi,k) \sin^2(\varphi) + F(\varphi,k) \cos^2(\varphi) \right) \tag{9}$$

where $\cos(\varphi) = \frac{c}{a}$, $k^2 = \frac{a^2(b^2-c^2)}{b^2(a^2-c^2)}$, and where $F(\varphi,k)$ and $E(\varphi,k)$ are incomplete elliptic integrals of the first and second kind respectively[5]. Keeping the longest semi-axis $a = a_0$ and solving the equation

$$S(a_0, b, c) = S(a_0, b_0, c_0) \tag{10}$$

with $a_0 = 4, b_0 = c_0 = 1$ by the commercial software WOLFRAM MATHEMATICA v.12.0.0.0, we can get $\beta := -\frac{b-b_0}{c-c_0} \approx -\left.\frac{\mathrm{d}b}{\mathrm{d}c}\right|_{b=c=1} = 1$ and the error in $\beta$ is less than 5% for the strain $\epsilon_b = \frac{b-b_0}{b} \leq 10\%$.

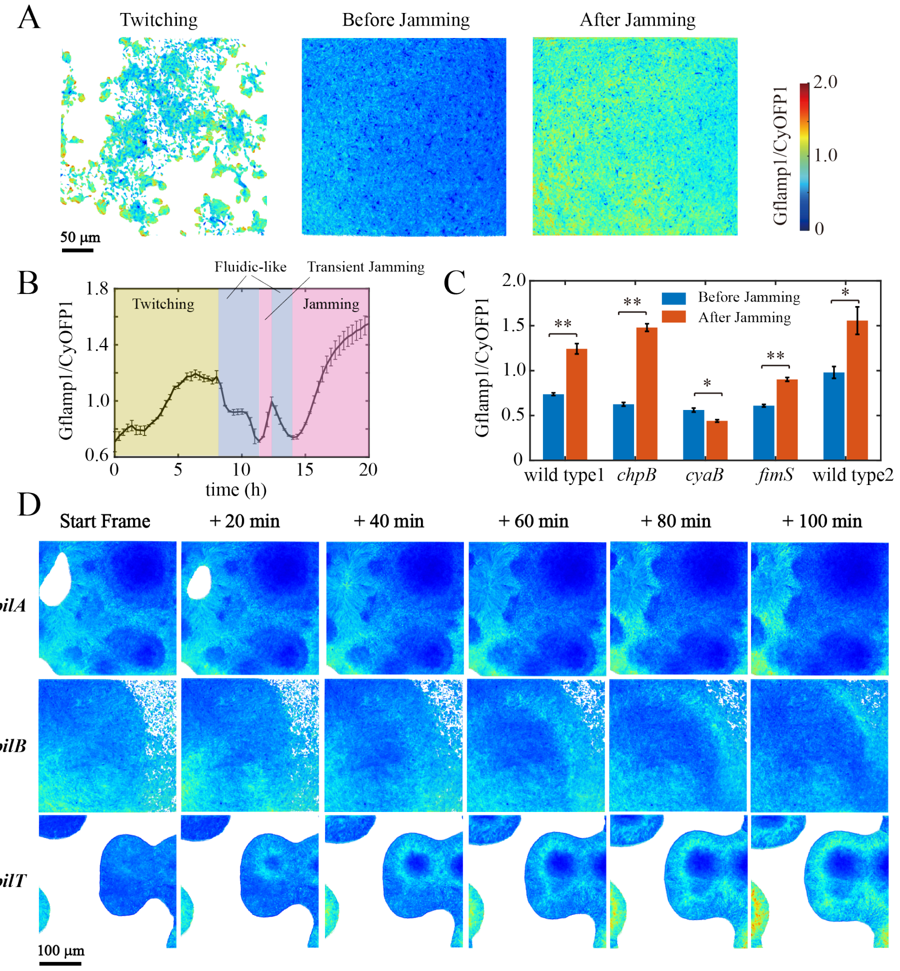


**Figure S1: Initial observations of Gflamp1/CyOFP1 signals of *P. aeruginosa* cells grown between hydrogel-glass interface.** (A) Spatial distributions of Gflamp1/CyOFP1 signal in wild-type P.*aeruginosa* during the stages of twitching motility, pre-jamming, and post-jamming phases at the hydrogel-glass interface. Scale bar, 50 µm. (B) Time series of mean Gflamp1/CyOFP1 signal in wild-type *P. aeruginosa* growing at the gel-glass interface. Yellow shaded area indicates the twitching motility phase, light purple shaded area denotes the fluidic-like phase, and magenta shaded area represents the jamming phase. Note the transient jamming periods during the fluidic-like phase, corresponding to short increases in Gflamp1/CyOFP1 signal. (C) Mean Gflamp1/CyOFP1 values 2 hours before and after entering the jamming phase for various mutant strains. Statistical significance determined by t-test, **, $p < 0.01$, *, $p < 0.05$. Wild type1, *cyaB*, and *chpB* using 1.5%(w/w) agarose, while *fimS* and wild type2 using 2.5%(w/w) agarose. (D) Spatial patterns of Gflamp1/CyOFP1 signal in twitching motility-deficient strains *pilA*, *pilB*, and *pilT* during growth at the gel-glass interface. Scale bar, 100 µm.

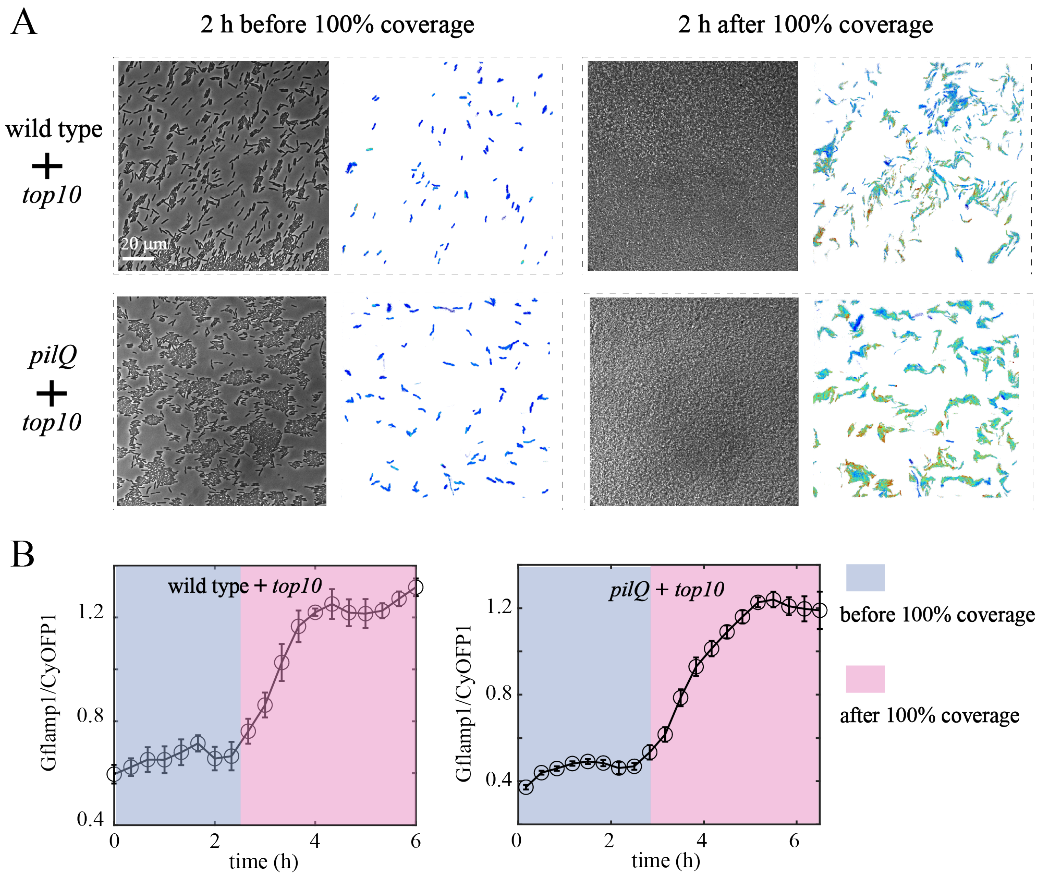


**Figure S2: Results of mixed culture experiments with *E. coli* and *P. aeruginosa* grown between hydrogel-glass interface.** (A) Phase contrast bright-field and Gflamp1/CyOFP1 fluorescence images of mixed cultures of *E. coli* TOP10 strain with wild-type or *pilQ* mutant *P. aeruginosa* strain under 2.5% (w/w) agarose gel. Images were taken 2 hours before and after bacterial colonization of the entire field of view. Scale bar, 20 µm. (B) Time-series of Gflamp1/CyOFP1 signal in mixed cultures experiments corresponding to data in (A). light purple shaded area denotes the phase before 100% field coverage, and magenta shaded area represents the phase after 100% field coverage.

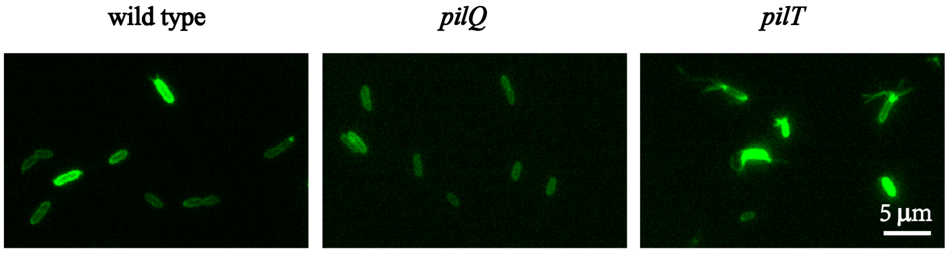


**Figure S3: Staining of Type IV pili in the wild type, *pilQ* and *pilT* mutant strains of *P. aeruginosa*.** In wild-type cells, type IV pili were observed protruding from the poles of some cells. The *pilQ* mutant cells exhibited reduced staining intensity and no visible surface pili. The *pilT* mutant cells displayed multiple type IV pili extending from cell poles and demonstrated the highest degree of cellular staining. Scale bar, 5 µm.

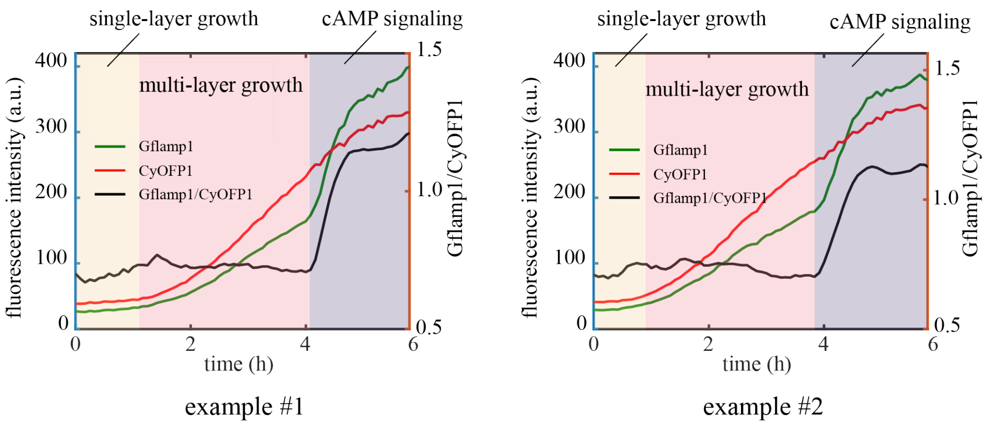


**Figure S4: Two experimental examples showing the temporal dynamics of Gflamp1 (green line), CyOFP1 fluorescence signals (red line), and the Gflamp1/CyOFP1 fluorescence ratio (black line) as *pilQ* mutant cells grow within gel microcavities.** The yellow region indicates single-layer bacterial growth, the pink region denotes transition into multi-layer growth, and the blue-purple region signifies the activation of cAMP signaling.

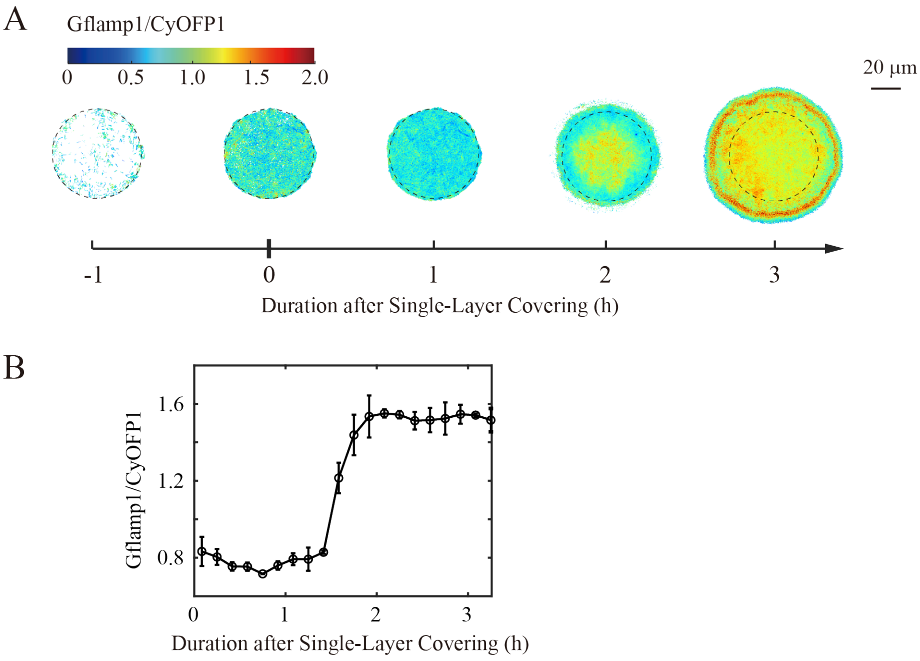


**Figure S5: Confined growth-induced compression triggers cAMP response in *P. aeruginosa*, 60 µm microcavities.** (A) Time-lapse ratio images of *P. aeruginosa pilQ* mutant strain grown in agarose microcavities (60 µm diameter, 2 µm depth). Scale bar: 20 µm. (B) Quantification of the Gflamp1/CyOFP1 ratio in relation to bacterial growth time after single-layer covering.

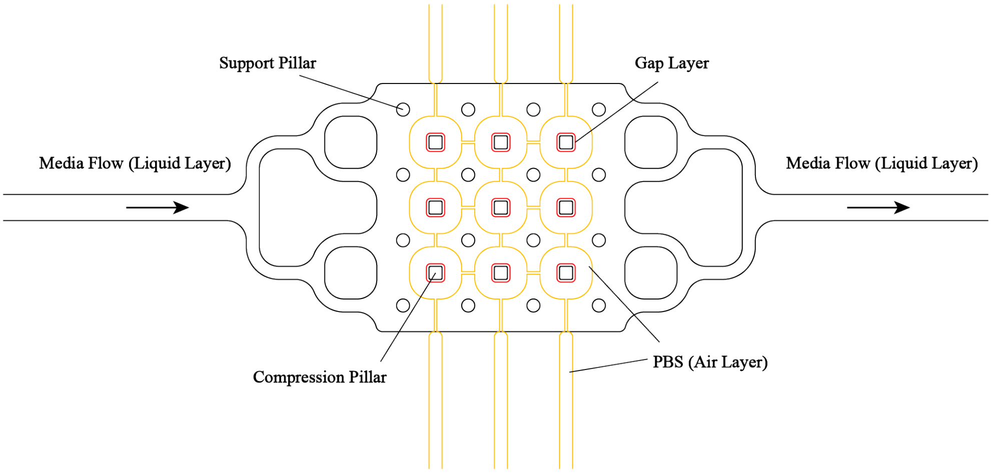


**Figure S6: Diagram showing the structure of the microfluidic device designed to apply mechanical pressure to bacterial cells.** The air layer (yellow lines) is filled with PBS buffer, and is 20 µm high and has 9 rounded square chambers connected by narrow channels. Each chamber is 400 µm in size. A thin PDMS membrane of 50 µm thickness seals the bottom of the air channels to create closed chambers for the air layer. The underside of the PDMS membrane and a glass slide form the liquid layer fluid channel, which is 18 µm high and allows bacterial cells to grow. The bottom of the PDMS membrane has two types of features (black lines): (1) nine square compression pillars for applying mechanical compression on bacteria, each 14.5 µm tall. These line up with the air-layer chambers above, so that the working air pressure in the air chamber will deform the PDMS membrane and make the pillars compress downward. (2) sixteen round support pillars, each 18 µm tall. The red square area (gap layer) in the diagram shows a raised part on the silicon mold used to make the PDMS membrane. This raised part is 3.5 µm high. It ensures that the compression-applying pillars on the PDMS membrane are shorter than the height of the fluid channel.

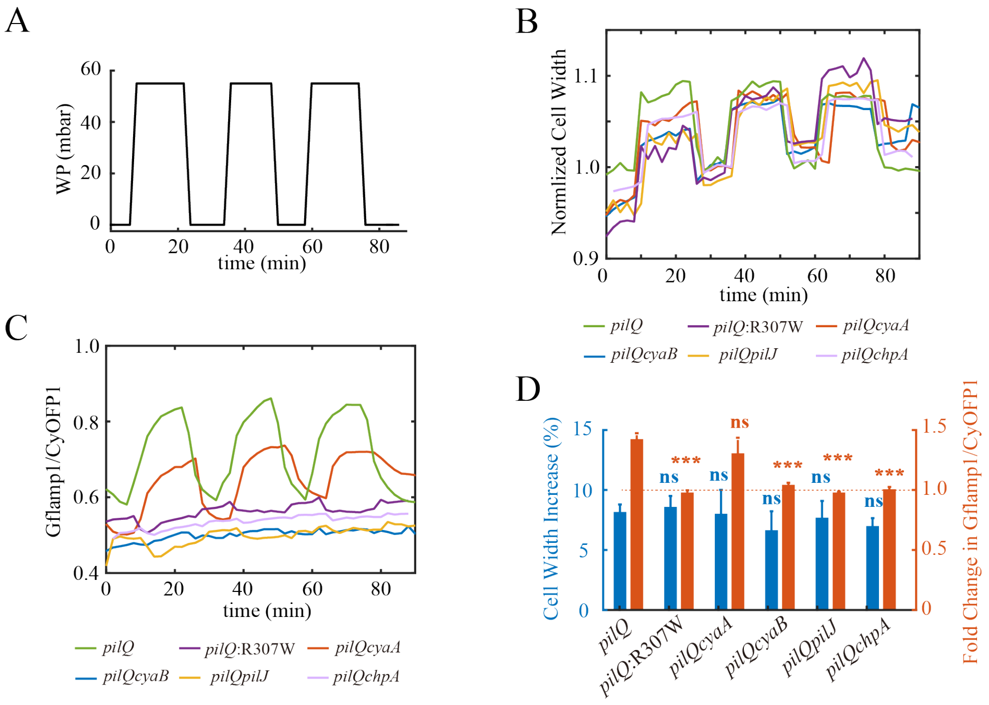


**Figure S7: Results of cyclic compression experiments on bacterial cells of various mutant strains derived from the *pilQ* knockout strain.** (A) Applied working pressure curve (identical to Fig. 2B). (B) Changes in average bacterial width in response to applied pressure. (C) Variations in mean Gflamp1/CyOFP1 signal intensity in response to applied pressure. (D) Percent change of bacterial width and fold change in Gflamp1/CyOFP1 under high working pressure in comparison to data under low working pressure. Statistical significance determined by t-test, ***, $p < 0.01$, ns, non significant.

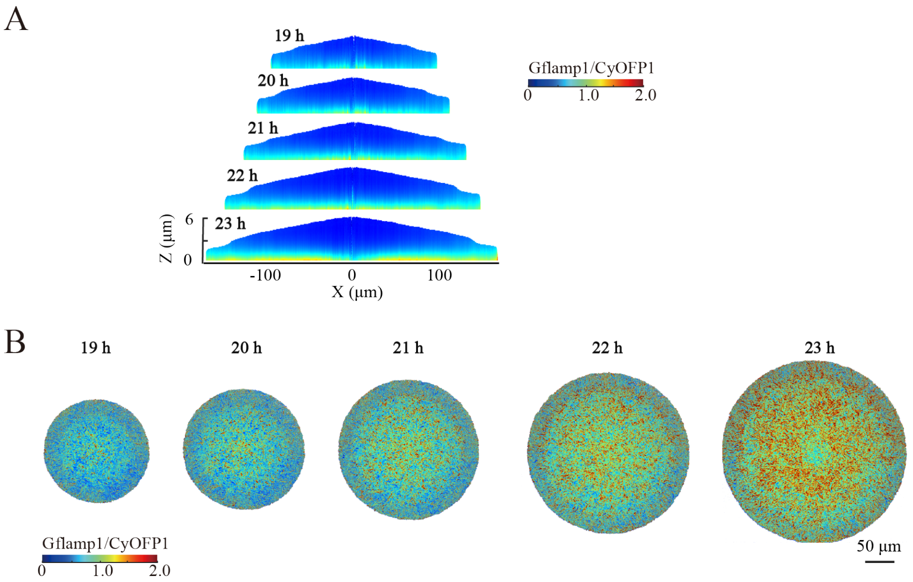


**Figure S8: The *pelpsl* mutant strain showed neither colony steps nor Gflamp1/CyOFP1 ring pattern.** (A) Z-direction scan images of colony cross-sections at different time points for a bacterial colony of the *pelBpslBCD* muant strain growing at the interface between 2.0% agarose gel and glass. (B) Time-lapse images showing the spatial distribution of Gflamp1/CyOFP1 signal in the bacterial layer directly adhering to the glass surface (corresponding to data in panel A). Scale bar, 50 μm.

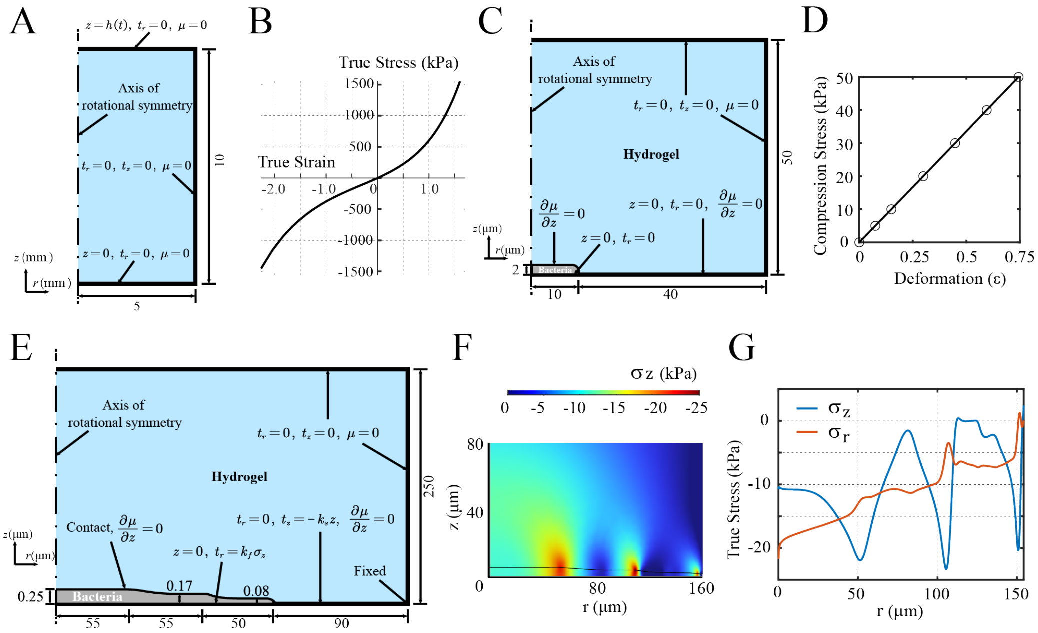


**Figure S9: Physical modeling and numerical simulation of forces acting on bacteria under gel compression.** (A) nitial geometry with Dirichlet and Neumann boundary conditions used in the calculation of uniaxial compression and stretch of cylindrical hydrogels. (B) Stress-strain curve of uniaxial test simulations with polymer mass fraction and the Flory–Huggins parameter. (C) Initial geometry with Dirichlet and Neumann boundary conditions used in the calculation of bacteria filling a cavity. The bottom of hydrogel is always attached to the substrate. (D) The relationship between the equilibrium stress in the $z-$direction and the final thickness of the bacteria with an initial thickness of 2 µm appears to be linear. (E) Initial geometry (zoomed in the $z-$direction) with Dirichlet and Neumann boundary conditions used in the calculation of bacteria steps. (F) The $z-$component of true stress field when the thickness of the bacteria at reaches 5 µm shows stress concentrations at the step edges. (G) The $r-$ and $z-$components of true stress in the bottom layer of the bacteria.

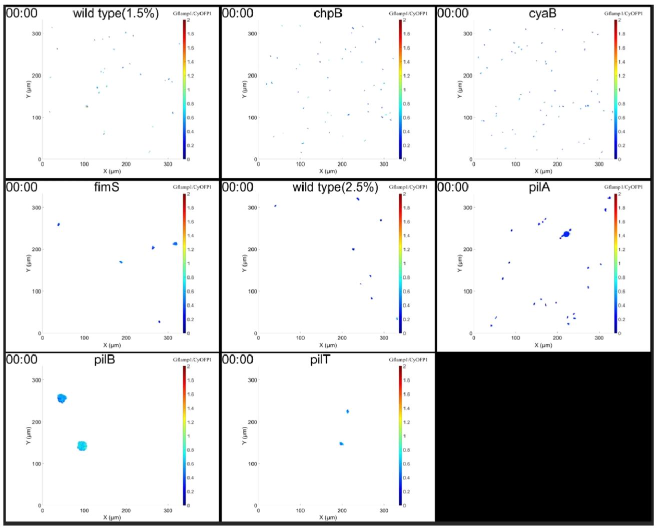


**Movie S1: Changes of Gflamp1/CyOFP1 signal in different P. aeruginosa mutant cells grown at hydrogel-glass interface.** Wild type (1.5%), *cyaB*, and *chpB* using 1.5%(w/w) agarose, while *fimS*, wild type (2.5%), *pilA*, *pilB*, and *pilT* using 2.5%(w/w) agarose.

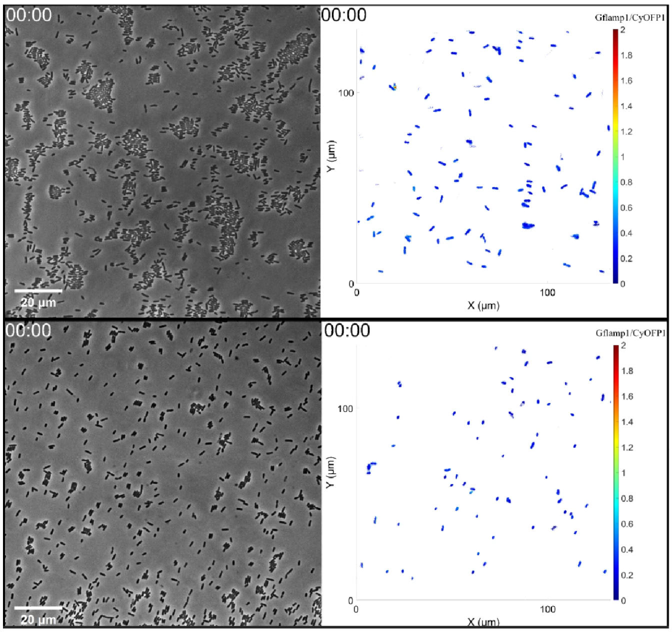


**Movie S2: Change of Gflamp1/CyOFP1 in P. aeruginosa cells in mixed culture experiments in which cells of top10 and wild type of *pilQ* mutant strain of P. aeruginosa were grown between hydrogel-glass interface.** Left, phase contrast bright field movies. Right, Gflamp1/CyOFP1 movies. Scale bar, 20 µm.

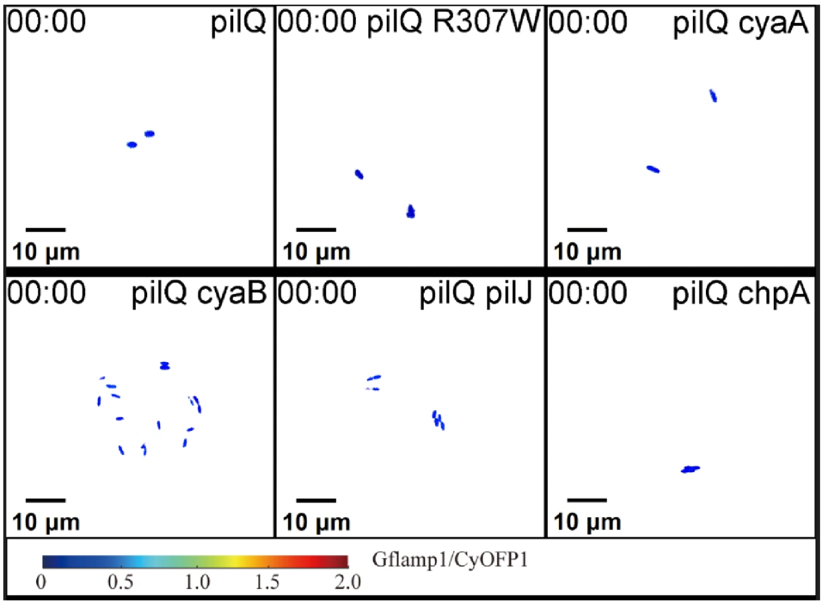


**Movie S3: Changes of Gflamp1/CyOFP1 signal in different *P. aeruginosa* mutant cells grown in agarose microcavities** (30 µm **diameter,** 2 µm **depth).** Scale bar: 10 µm.

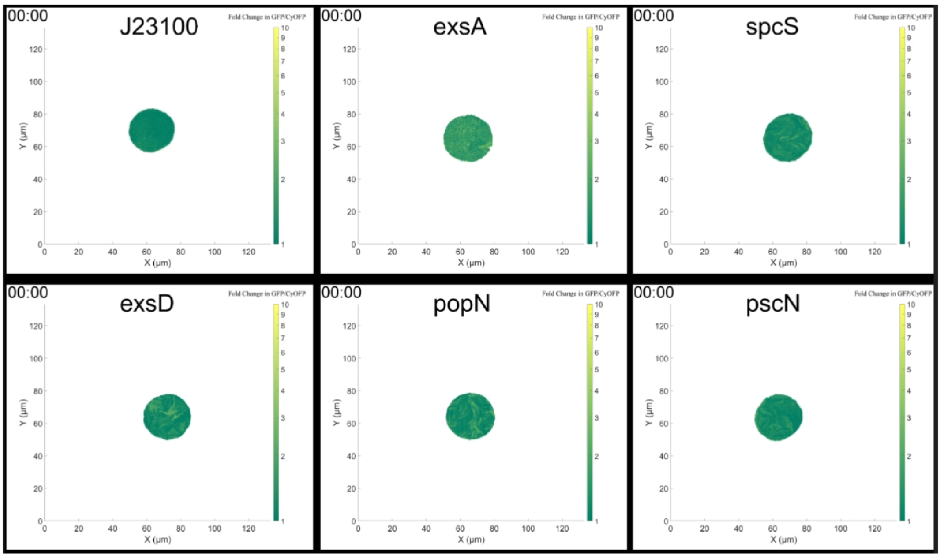


**Movie S4: Changes of SfGFP/CyOFP1 signal in different transcriptional reporters expressed in the *pilQ* mutant strain grown in agarose microcavities (30 µm diameter, 2 µm depth).**

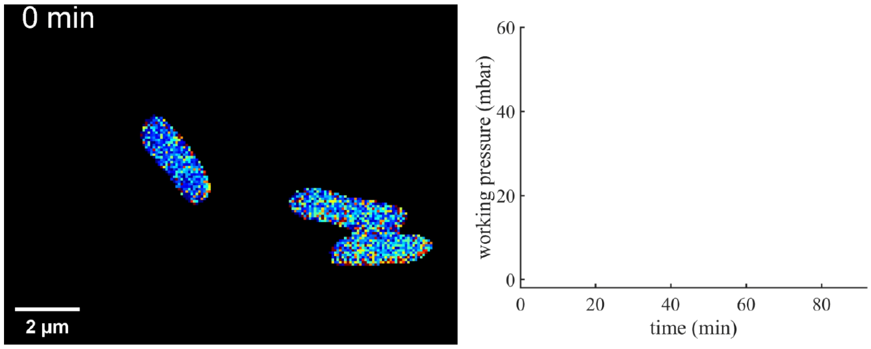


**Movie S5: A typical example of intracellular cAMP response to cyclic compression pressure in *pilQ* mutant cells.** The left panel shows Gflamp1/CyOFP1 signal, the right panel displays the corresponding applied pressure over time.

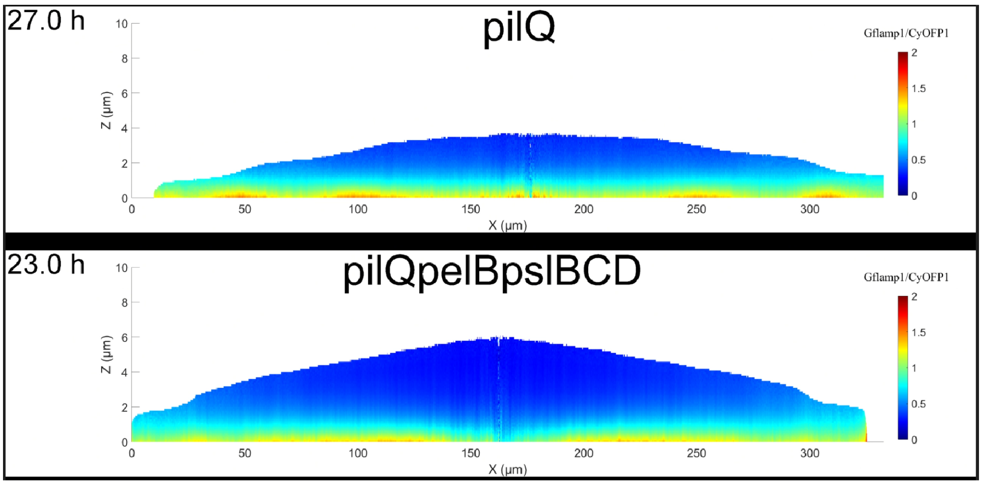


**Movie S6: Movies of colony cross-sections at different time points for *pilQ* and *pilQpelBpslBCD* strains grown at the interface between 2.0% agarose gel and glass**

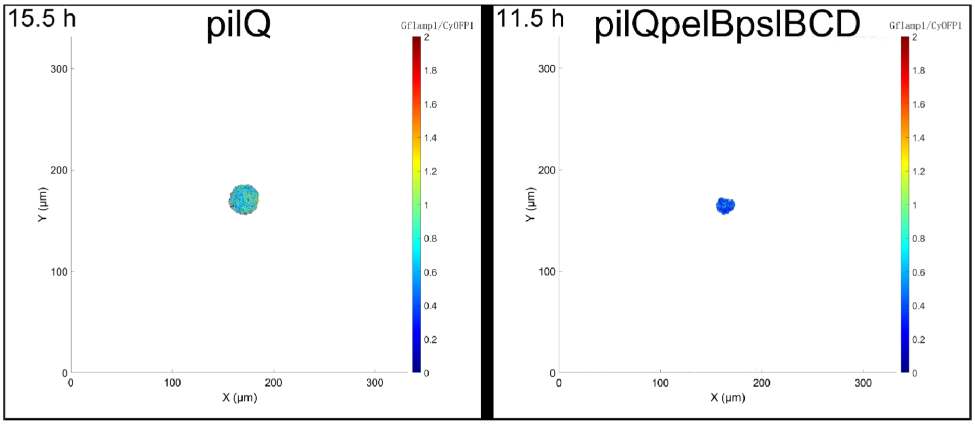


**Movie S7: Movies showing the spatial distribution of Gflamp1/CyOFP1 signals in the bacterial layer directly adhering to the glass surface.** The concentration of agarose hydrogel is 2% (w/w). Left, *pilQ* colony. Right, *pilQpelBpslBCD* colony.

#### Table S1. Strains and Plasmids used in this study

**Table 1**: Strains and Plasmids used in this study

| Strains or plasmids | Description | Origin |
|---|---|---|
| ***E. coli* strain** | | |
| Top10 | $F^-$, *mcrA*, (*mrr, hsdRMS–mcrBC*), $\phi$80*lacZ* M15 *lacX74, recA*1, *araD*139, (Δara–leu)7697, *galU, galK, rpsL*(Str$^R$), *endA1, nupG* | Invitrogen |
| ***P. aeruginosa* strains** | | |
| PAO1 | Wild-type strain | J.D. Shrout |
| PAO1-Δ*pilQ* | nonpolar *pilQ* deletion in PAO1 | This study |
| PAO1-Δ*cyaA* | nonpolar *cyaA* deletion in PAO1 | This study |
| PAO1-Δ*cyaB* | nonpolar *cyaB* deletion in PAO1 | This study |
| PAO1-Δ*pilJ* | nonpolar *pilJ* deletion in PAO1 | This study |
| PAO1-Δ*chpA* | nonpolar *chpA* deletion in PAO1 | This study |
| PAO1-Δ*pilT* | nonpolar *pilT* deletion in PAO1 | This study |
| PAO1-Δ*pilQ*Δ*cyaA* | nonpolar *pilQ* and *cyaA* deletion in PAO1 | This study |
| PAO1-Δ*pilQ*Δ*cyaB* | nonpolar *pilQ* and *cyaB* deletion in PAO1 | This study |
| PAO1-Δ*pilQ*Δ*pilJ* | nonpolar *pilQ* and *pilJ* deletion in PAO1 | This study |
| PAO1-Δ*pilQ*Δ*chpA* | nonpolar *pilQ* and *chpA* deletion in PAO1 | This study |
| PAO1-pJN105-J23100-cyOFP1-gly4-gflamp1 | WT strain containing *J23100-cyOFP1-gly4-gflamp1*-pJN105, Gflamp1 expression under the constitute promoter *J23100*, Gm$^R$ | This study |

Continued on next page

Table S1 continued

| **Strains or plasmids** | **Description** | **Origin** |
|---|---|---|
| PAO1-Δ*pilQ*-pJN105-J23100-cyOFP1-gly4-gflamp1 | PAO1-Δ*pilQ* strain containing *J23100-cyOFP1-gly4-gflamp1*-pJN105, Gflamp1 expression under the constitute promoter *J23100*, $Gm^R$ | This study |
| PAO1-Δ*pilQ*Δ*cyaA*-pJN105-J23100-cyOFP1-gly4-gflamp1 | PAO1-Δ*pilQ*Δ*cyaA* strain containing *J23100-cyOFP1-gly4-gflamp1*-pJN105, Gflamp1 expression under the constitute promoter *J23100*, $Gm^R$ | This study |
| PAO1-Δ*pilQ*Δ*cyaB*-pJN105-J23100-cyOFP1-gly4-gflamp1 | PAO1-Δ*pilQ*Δ*cyaB* strain containing *J23100-cyOFP1-gly4-gflamp1*-pJN105, Gflamp1 expression under the constitute promoter *J23100*, $Gm^R$ | This study |
| PAO1-Δ*pilQ*Δ*pilJ*-pJN105-J23100-cyOFP1-gly4-gflamp1 | PAO1-Δ*pilQ*Δ*pilJ* strain containing *J23100-cyOFP1-gly4-gflamp1*-pJN105, Gflamp1 expression under the constitute promoter *J23100*, $Gm^R$ | This study |
| PAO1-Δ*pilQ*Δ*chpA*-pJN105-J23100-cyOFP1-gly4-gflamp1 | PAO1-Δ*pilQ*Δ*chpA* strain containing *J23100-cyOFP1-gly4-gflamp1*-pJN105, Gflamp1 expression under the constitute promoter *J23100*, $Gm^R$ | This study |
| PAO1-Δ*pilQ*-pJN105-J23100-cyOFP1-gly4-gflamp1-mutR307W | PAO1-Δ*pilQ* strain containing *J23100-cyOFP1-gly4-gflamp1*-pJN105, Gflamp1 R307W mutant expression under the constitute promoter *J23100*, $Gm^R$ | This study |

Continued on next page

Table S1 continued

| **Strains or plasmids** | **Description** | **Origin** |
|---|---|---|
| PAO1-Δ*pilQ*-pJN105-P*popN*-sfGFP-T-cyOFP1 | PAO1-Δ*pilQ* strain containing a reporter plasmid, SfGFP expression under the promoter of P*popN*, $Gm^R$ | This study |
| PAO1-Δ*pilQ*-pJN105-P*exoS*-sfGFP-T-cyOFP1 | PAO1-Δ*pilQ* strain containing a reporter plasmid, SfGFP expression under the promoter of P*exoS*, $Gm^R$ | This study |
| PAO1-Δ*pilQ*-pJN105-P*exoT*-sfGFP-T-cyOFP1 | PAO1-Δ*pilQ* strain containing a reporter plasmid, SfGFP expression under the promoter of P*exoT*, $Gm^R$ | This study |
| PAO1-Δ*pilQ*-pJN105-P*spcS*-sfGFP-T-cyOFP1 | PAO1-Δ*pilQ* strain containing a reporter plasmid, SfGFP expression under the promoter of P*spcS*, $Gm^R$ | This study |
| PAO1-Δ*pilQ*-pJN105-P*exsC*-sfGFP-T-cyOFP1 | PAO1-Δ*pilQ* strain containing a reporter plasmid, SfGFP expression under the promoter of P*exsC*, $Gm^R$ | This study |
| PAO1-Δ*pilQ*-pJN105-P*exsA*-sfGFP-T-cyOFP1 | PAO1-Δ*pilQ* strain containing a reporter plasmid, SfGFP expression under the promoter of P*exsA*, $Gm^R$ | This study |
| PAO1-Δ*pilQ*-pJN105-P*exsD*-sfGFP-T-cyOFP1 | PAO1-Δ*pilQ* strain containing a reporter plasmid, SfGFP expression under the promoter of P*exsD*, $Gm^R$ | This study |
| PAO1-Δ*pilQ*-pJN105-P*pscN*-sfGFP-T-cyOFP1 | PAO1-Δ*pilQ* strain containing a reporter plasmid, SfGFP expression under the promoter of P*pscN*, $Gm^R$ | This study |

Continued on next page

Table S1 continued

| **Strains or plasmids** | **Description** | **Origin** |
|---|---|---|
| PAO1-Δ*pilQ*-pJN105-*J23100*-sfGFP-T-cyOFP1 | PAO1-Δ*pilQ* strain containing a reporter plasmid, SfGFP expression under the promoter of *J23100*, $Gm^R$ | This study |
| PAO1-Δ*pilQ*Δ*pilJ*-pJN105-P*popN*-sfGFP-T-cyOFP1 | PAO1-Δ*pilQ*Δ*pilJ* strain containing a reporter plasmid, SfGFP expression under the promoter of P*popN*, $Gm^R$ | This study |
| PAO1-Δ*pilQ*Δ*pilJ*-pJN105-P*exoS*-sfGFP-T-cyOFP1 | PAO1-Δ*pilQ*Δ*pilJ* strain containing a reporter plasmid, SfGFP expression under the promoter of P*exoS*, $Gm^R$ | This study |
| PAO1-Δ*pilQ*Δ*pilJ*-pJN105-P*exoT*-sfGFP-T-cyOFP1 | PAO1-Δ*pilQ*Δ*pilJ* strain containing a reporter plasmid, SfGFP expression under the promoter of P*exoT*, $Gm^R$ | This study |
| PAO1-Δ*pilQ*Δ*pilJ*-pJN105-P*spcS*-sfGFP-T-cyOFP1 | PAO1-Δ*pilQ*Δ*pilJ* strain containing a reporter plasmid, SfGFP expression under the promoter of P*spcS*, $Gm^R$ | This study |
| PAO1-Δ*pilQ*Δ*pilJ*-pJN105-P*exsC*-sfGFP-T-cyOFP1 | PAO1-Δ*pilQ*Δ*pilJ* strain containing a reporter plasmid, SfGFP expression under the promoter of P*exsC*, $Gm^R$ | This study |
| PAO1-Δ*pilQ*Δ*pilJ*-pJN105-P*exsA*-sfGFP-T-cyOFP1 | PAO1-Δ*pilQ*Δ*pilJ* strain containing a reporter plasmid, SfGFP expression under the promoter of P*exsA*, $Gm^R$ | This study |
| PAO1-Δ*pilQ*Δ*pilJ*-pJN105-P*exsD*-sfGFP-T-cyOFP1 | PAO1-Δ*pilQ*Δ*pilJ* strain containing a reporter plasmid, SfGFP expression under the promoter of P*exsD*, $Gm^R$ | This study |

Continued on next page

Table S1 continued

| **Strains or plasmids** | **Description** | **Origin** |
|---|---|---|
| PAO1-Δ*pilQ*Δ*pilJ*-pJN105-P*pscN*-sfGFP-T-cyOFP1 | PAO1-Δ*pilQ*Δ*pilJ* strain containing a reporter plasmid, SfGFP expression under the promoter of P*pscN*, $Gm^R$ | This study |
| PAO1-Δ*pilQ*Δ*pilJ*-pJN105-*J23100*-sfGFP-T-cyOFP1 | PAO1-Δ*pilQ*Δ*pilJ* strain containing a reporter plasmid, SfGFP expression under the promoter of *J23100*, $Gm^R$ | This study |
| PAO1-pucp20-Pbad-*pilA*-S99C-T-J23100-*rflamp* | PAO1 strain containing a plasmid for staining of type IV pili, $Gm^R$ | This study |
| PAO1-Δ*pilQ*-pucp20-Pbad-*pilA*-S99C-T-J23100-*rflamp* | PAO1-Δ*pilQ* strain containing a plasmid for staining of type IV pili, $Gm^R$ | This study |
| PAO1-Δ*pilT*-pucp20-Pbad-*pilA*-S99C-T-J23100-*rflamp* | PAO1-Δ*pilT* strain containing a plasmid for staining of type IV pili, $Gm^R$ | This study |
| **Plasmids** | | |
| pex18gm | $oriT^+$ $sacB^+$; gene replacement vector with MCS from pUC18; $Gm^R$ | This study |
| *J23100-cyOFP1-gly4-gflamp1*-pJN105 | Expression vector with Gflamp1 protein for indicating the internal cAMP level | This study |
| *J23100-cyOFP1-gly4-gflamp1-mutR307W*-pJN105 | Expression vector with Gflamp1 R307W mutant protein as control | This study |

Continued on next page

Table S1 continued

| Strains or plasmids | Description | Origin |
|---|---|---|
| pJNTR | Intermediate plasmid with the gene expression module RBSII-sfGFP-T0T1-J23102-CyOFP1 for further construction of transcriptional reporter, $Gm^R$ | This study |
| P*popN*-sfGFP-T-cyOFP1-pJN105 | Transcriptional reporter plasmid of *popN*, $Gm^R$ | This study |
| P*exoS*-sfGFP-T-cyOFP1-pJN105 | Transcriptional reporter plasmid of *exoS*, $Gm^R$ | This study |
| P*exoT*-sfGFP-T-cyOFP1-pJN105 | Transcriptional reporter plasmid of *exoT*, $Gm^R$ | This study |
| P*spcS*-sfGFP-T-cyOFP1-pJN105 | Transcriptional reporter plasmid of *spcS*, $Gm^R$ | This study |
| P*exsC*-sfGFP-T-cyOFP1-pJN105 | Transcriptional reporter plasmid of *exsC*, $Gm^R$ | This study |
| P*exsA*-sfGFP-T-cyOFP1-pJN105 | Transcriptional reporter plasmid of *exsA*, $Gm^R$ | This study |
| P*pscN*-sfGFP-T-cyOFP1-pJN105 | Transcriptional reporter plasmid of *pscN*, $Gm^R$ | This study |
| P*exsD*-sfGFP-T-cyOFP1-pJN105 | Transcriptional reporter plasmid of *exsD*, $Gm^R$ | This study |
| *J23100*-sfGFP-T-cyOFP1-pJN105 | Transcriptional reporter plasmid of *J23100* as control, $Gm^R$ | This study |
| pCasPA | Modified plasmid for in-frame deletion via CRISPR system from pCas; $tet^R$ | This study |

Continued on next page

Table S1 continued

| **Strains or plasmids** | **Description** | **Origin** |
| --- | --- | --- |
| pACRISPR | Modified plasmid with sgRNA sequence and homologous sequences for in-frame deletion; $Ap^R$ | This study |
| *cyaA*-pex18gm | In-frame deletion of *cyaA* in pex18gm; $Gm^R$ | This study |
| *cyaB*-pex18gm | In-frame deletion of *cyaB* in pex18gm; $Gm^R$ | This study |
| *pilQ*-pACRISPR | In-frame deletion of *pilQ* in plasmid pACRISPR; $Ap^R$ | This study |
| *chpA*-pACRISPR | In-frame deletion of *chpA* in plasmid pACRISPR; $Ap^R$ | This study |
| *pilJ*-pACRISPR | In-frame deletion of *pilJ* in plasmid pACRISPR; $Ap^R$ | This study |
| Pbad-*pilA*-S99C-T-J23100-*rflamp*-pucp20 | Expression vector with S99C mutant in *pilA* for staining of type IV pili, $Gm^R$ | This study |

#### Table S2. Primers used in this study

**Table 2**: Primers used in this study

| Primer name | Oligonucleotide Sequence |
|---|---|
| *pJN105 cloning primers* | |
| pJN105-F | GCGCCTAAcccgtttttttgggctagcgaattcctgcagcc |
| pJN105-R | tagcactgtacctaggactgagctagccgtcaatatttgccaata |
| J23100-F | tagctcagtcctaggtacagtgcta |
| Gflamp-R | ccaaaaaaacgggTTAGGCGC |
| *Gene knockout primers* | |
| PRISPOR-F | ctcgagacttttcatactcccgccattc |
| N20-R | tctagaccatgggtatggacagatctc |
| chpA-N20-R | aaaacCTGGACGATTCCGCCGAAGTccacacattatacgagccggatgattaattgtca |
| chpA-N20-F | ACTTCGGCGGAATCGTCCAGgttttagagctagaaatagcaagttaaaataaggctagt |
| chpA-up-F | gatctgtccatacccatggTCTAGAAACCCTACTCGCTGGCGATG |
| chpA-up-R | GTCACCCATAGCCACTCCATT |
| chpA-dn-F | CATGAATGGAGTGGCTATGGGTGACGAAGCCATCCAGTCCCTGGT |
| chpA-dn-R | tggcgggagtatgaaaagtCTCGAGGCGTCGAGAAATGCCTTCAC |
| pilQ-N20-F | TCGAGGCGAAGGATCGCACAgttttagagctagaaatagcaagttaaaataaggctagt |
| pilQ-N20-R | TGTGCGATCCTTCGCCTCGAccacacattatacgagccggatgattaattgtca |

Continued on next page

Table S2 continued

| Primer name | Oligonucleotide Sequence |
|---|---|
| pilQ-up-F | gatctgtccatacccatggTCTAGAGAATTTCCTGGACGACCAGG |
| pilQ-up-R | ATCGCAATCGGTCGCTGATA |
| pilQ-dn-F | TATCAGCGACCGATTGCGATACTGTTCATCGTCCGACTCC |
| pilQ-dn-R | tggcgggagtatgaaaagtCTCGAGTCCTACATGGACGAGGTTCG |
| pilJ-N20-F | TCGGTAGCCATTGCCAACAAgttttagagctagaaatagcaagttaaaataaggctagt |
| pilJ-N20-R | aaaacTTGTTGGCAATGGCTACCGAccacacattatacgagccggatgattaattgtca |
| pilJ-up-F | gagatctgtccatacccatggTCTAGAGGTTCTTCGTCGCCCCCATG |
| pilJ-up-R | GCCTGCGTTGATTTTCTTCA |
| pilJ-dn-F | TGAAGAAAATCAACGCAGGCGTATCCGGCTTCAAACTGCC |
| pilJ-dn-R | gaatggcgggagtatgaaaagtCTCGAGGCCAGCGAGTAGGGTTCCTC |
| *Primers for transcription* | |
| VecGFPOFP-F | TTTCACAAAAGAGGAGAAAggcatt |
| VecGFPOFP-R | caatatttgccaatacggggttctt |
| *popN*-F | aagaaccccgtattggcaaatattgGAGCAGGTGCTCTCCGACCG |
| *popN*-R | aatgccTTTCTCCTCTTTTGTGAAATCCTGGTCTGCAAAGGCGGC |

Continued on next page

Table S2 continued

| **Primer name** | **Oligonucleotide Sequence** |
|---|---|
| *exoS*-F | aagaaccccgtattggcaaatattgGTTGTTCGAGTTGATGGTGG |
| *exoS*-R | aatgccTTTCTCCTCTTTTGTGAAACTCCTGATGTTTC TCCGCCA |
| *exoT*-F | aagaaccccgtattggcaaatattgCAGGCTGAAGGTGCGGATTC |
| *exoT*-R | aatgccTTTCTCCTCTTTTGTGAAACTCCTGATGTTTC CCCGCCA |
| *exsC*-F | aagaaccccgtattggcaaatattgAAGATGCAGGCGCTTGGCAA |
| *exsC*-R | aatgccTTTCTCCTCTTTTGTGAAATAAAGCTCAGCGC ATGCTAG |
| *exsD*-F | aagaaccccgtattggcaaatattgCCTGAAGATCGAGGAGTTGC |
| *exsD*-R | aatgccTTTCTCCTCTTTTGTGAAACTTCCTCACTACAT TGCGCG |
| *pscN*-F | aagaaccccgtattggcaaatattgCCCTCCAGGTAAGCGCTGAG |
| *pscN*-R | aatgccTTTCTCCTCTTTTGTGAAACGGCTGGAGTCGG CGCGCACTATA |
| *exsA*-F | aagaaccccgtattggcaaatattgTTGCCGTTGCGCTATGCCTT |
| *exsA*-R | aatgccTTTCTCCTCTTTTGTGAAACCAACACTTCCCG TCGTACC |
| *spcs*-F | aagaaccccgtattggcaaatattgAGCACCATCTGCTTGAACAC |
| *spcs*-R | aatgccTTTCTCCTCTTTTGTGAAAGTCACTGGAGGCA GCCATTA |
